# KDM6B sensitizes PARthanatos via suppression of O^6^-methylguanine DNA methyltransferase repair and sustained checkpoint response

**DOI:** 10.1101/2021.12.17.473255

**Authors:** Mingming Yang, Chenliang Wang, Mi Zhou, Lei Bao, Yanan Wang, Ashwani Kumar, Chao Xing, Weibo Luo, Yingfei Wang

## Abstract

Poly(ADP-ribose) polymerase-1 (PARP-1) is a DNA damage sensor and contributes to both DNA repair and cell death processes. However, how PARP-1 signaling is regulated to switch its function from DNA repair to cell death remains largely unknown. Here, we found that PARP-1 plays a central role in alkylating agent-induced PARthanatic cancer cell death. Lysine demethylase 6B (KDM6B) was identified as a key cell death effector in PARthanatos. Knockout of KDM6B or loss of KDM6B demethylase activity conferred cancer cells resistance to PARthanatic cell death in response to alkylating agents. Mechanistically, KDM6B knockout suppressed methylation at the promoter of O^6^-methylguanine-DNA methyltransferase (MGMT) to enhance MGMT expression and its direct DNA repair function, thereby inhibiting DNA damage-evoked PARP-1 hyperactivation and subsequent cell death. Moreover, KDM6B knockout triggered sustained Chk1 phosphorylation and activated a second repair machinery to fix DNA damage evading from MGMT repair. Inhibition of MGMT or checkpoint response re-sensitized KDM6B deficient cells to PARthanatos induced by alkylating agents. These findings provide new molecular insights into epigenetic regulation of PARP-1 signaling mediating DNA repair or cell death and identify KDM6B as a biomarker for prediction of cancer cell vulnerability to alkylating agent treatment.

## Introduction

Poly(ADP-ribose) polymerase-1 (PARP-1) is a ubiquitously expressed nuclear enzyme and plays a critical role in DNA damage response in human cancers (1–3). Upon DNA damage, PARP-1 senses DNA single-strand breaks and utilizes NAD^+^ as the substrate to catalyze the addition of poly(ADP-ribose) (PAR) toward different acceptor proteins, including PARP-1 itself (1–3). As a result, PARP-1 activation leads to the recruitment of DNA repair proteins and nucleases to sites of DNA damage, thereby facilitating DNA damage repair (4). Inhibition of PARP-1 impairs DNA single-strand break repair and eventually causes DNA double-strand breaks, which are primarily repaired through homologous recombination and/or non-homologous end joining mechanisms (5). Blockade of DNA double-strand break repair in BRCA-deficient cancer cells can synergize with PARP inhibitor to induce cancer cell death (6). Accumulating studies reveal a dual function of PARP-1 in cell death and survival. PARP-1 hyperactivation promotes cell death, also known as PARP-1-dependent cell death (PARthanatos), in neurological diseases (7). After PARP-1 is activated by oxidative stress under neuronal injury, PAR functions as a death signal triggering apoptosis-inducing factor (AIF) translocation from mitochondria to the nucleus, where it interacts with the nuclease macrophage migration inhibitory factor (MIF) leading to large DNA fragmentation and neuronal cell death (8,9). However, whether PARthanatos also occurs in cancer cells and how PARP-1 signaling is regulated and switched between DNA repair and cell death remain poorly understood.

Alkylating agents are the most common and oldest chemotherapy drugs currently used in the clinic to kill cancer (10–12). Alkylating agents are divided into five categories: nitrogen mustard, nitrosoureas, alkyl sulfonates, triazines, and ethylenimines. *N*-Methyl-*N*′-nitro-*N*-nitrosoguanidine (MNNG), belonging to the nitrosourea category, was first discovered in 1950s as an anti-cancer drug through a national cooperative cancer drug screen (12). MNNG contains the most basic structure similar to several commonly used alkylating chemotherapy drugs Temozolomide (TMZ), Streptozotocin, Lomustine and Carmustine (10). It acts by producing an intermediate product methyldiazonium ion and adds alkyl groups specifically to the O^6^ of guanine and O^4^ of thymine, leading to DNA damage and cell death (12). Previous studies identified that mitochondrial dysfunction and NAD^+^ depletion contribute to alkylating agent-induced cell death (13,14). Human AlkB homolog 7 (ALKBH7) promotes loss of mitochondrial function and energy depletion in response to alkylating agents, leading to cell death (14). DNA damage repair proteins including O6-methylguanine-DNA methyltransferase (MGMT), exonuclease 1, and DNA mismatch repair (MMR) protein MutS Homolog 6 (MSH6) also play an important role in alkylating agent-induced cell death (15). Alkylating agents induce PARP-1 activation, but an early report suggest that PARP-1 activation is not sufficient to mediate cell death (13). Interestingly, the alkyladenine DNA glycosylase (MPG) is involved in alkylating agent-induced tissue damage *in vivo*, which is prevented by PARP-1 loss (16). Together, the precise role of PARP-1 in alkylating agent-induced cancer cell death and its signaling regulation are far not clear.

In the present study, we showed that PARP-1 is required for alkylating agents-induced PARthanatic cancer cell death and identified lysine demethylase 6B (KDM6B) as a key cell death effector in PARthanatos. KDM6B increased DNA damage by repressing both MGMT-mediated direct DNA repair and checkpoint response associated DNA repair, leading to PARP-1 hyperactivation and subsequent PARthanatic cell death. Inhibition of MGMT or checkpoint response sensitized PARthanatos induced by alkylating agents. These findings unravel an epigenetic mechanism that controls PARP-1-dependent DNA repair *vs*. cell death pathways and cancer cell vulnerability to alkylating agents.

## Materials and Methods

### Plasmid constructs

Full-length and C-terminal truncated KDM6B cDNA was amplified by PCR from KDM6B plasmids (Addgene, #21212 and #21214) and subcloned into pLVX-Ubc-FLAG vector. Full-length MGMT cDNA was amplified by PCR and cloned into cFUGW-3xflag-N vector. Catalytically inactive MGMT mutant was generated by site-directed mutagenesis PCR. DNA oligonucleotides of the single guide RNA (sgRNA) targeting human KDM6B, PARP-1, or MGMT (Supplementary Table S1) were annealed and ligated into BsmBI-linearized lentiCRISPRv2 vector (Addgene, #52961). Other plasmids have been described previously (8,17). All recombinant plasmids were verified by Sanger sequencing.

### Cell culture

HeLa, HEK293T, MDA-MB-231, MCF-7, and SUM159 cells were cultured in DMEM or DMEM/Ham’s F-12 supplemented with 10% heat-inactivated fetal bovine serum at 37°C in a 5% CO_2_/95% air incubator. All KO cell lines were generated using the CRISPR/Cas9 technique and genotyped as described (18). All cell lines were annually tested to be mycoplasma-free and authenticated by STR DNA profiling analysis during 2016-2017.

### Clonogenic assay

300 cells/well were seeded on a 24-well plate and treated with indicated drugs for 10 days. Colonies were washed with phosphate-buffered saline (PBS), fixed with 100% methanol, and stained with 0.01% crystal violet. Colony numbers were counted for quantification.

### Cell death assay

2×10^5^ cells/well were seeded on a 12-well plate and pretreated with vehicle, cell death inhibitors, PARP inhibitor DPQ (30 μM), GDC0575 (50 nM), or BG (200 μM) for 30 min followed by MNNG (50 μM) treatment for 15 min or methyl methanesulfonate (MMS, 2 mM) for 1 h in the presence or absence of indicated inhibitors. After the treatment with alkylating agents, cells were changed to fresh medium with or without indicated inhibitors for continuous incubation for 24-96 h. Cells were stained with 2 μM propidium iodide (PI) and 7 μM Hoechst 33342 (ThermoFisher) for 5-10 min and imaged under a Zeiss Observer Z1 microscope. The numbers of total and PI-positive cells were quantified as described (8,9).

### CRISPR screening and bioinformatics analysis

HeLa cells were transduced overnight at a MOI of 0.4 with a pooled genome-wide CRISPR KO (GeCKO v2) library A or B (Addgene) containing a total of 122,411 sgRNAs (6 sgRNAs per gene, 4 sgRNAs per miRNA, and 1000 control sgRNAs). After puromycin (1 μg/ml) selection, cells were treated with MNNG (75 μM) for 15 min, followed by continuous incubation with fresh medium for 3 days. The majority of cells died 24 h after treatment. The survived cells were pooled and subjected to genomic DNA extraction. The sgRNA library was amplified by two steps of PCR as described previously (19). Samples were sequenced on Illumina NextSeq 500 with read configuration as 150 bp, single end. The fastq files were subjected to quality check using fastqc (version 0.11.2, http://www.bioinformatics.babraham.ac.uk/projects/fastqc) and fastq_screen (version 0.4.4, http://www.bioinformatics.babraham.ac.uk/projects/fastq_screen), and adapters trimmed using an in-house script. The reference sgRNA sequences for human GeCKO v2.0 (A and B) were downloaded from Addgene (https://www.addgene.org/pooled-library/). The trimmed fastq files were mapped to reference sgRNA library using MAGeCK (20). Further, read counts for each sgRNA were generated and median normalization was performed to adjust for library sizes. Positively and negatively selected sgRNA and genes were identified using the default parameters of MAGeCK.

### RNA-Seq

RNA-Seq was performed as described previously (17). Total RNA was isolated from parental, KDM6B KO2, and KDM6B rescue HeLa cells using the RNeasy mini kit (Qiagen) and treated with DNase (Qiagen). The quality of total RNA was confirmed with a RNA integrity number score 8.5 or higher by the Agilent Tapestation 4200. mRNAs were used for library preparation with a Kapa mRNA HyperPrep kit (Roche) and sequenced on the Illumina NextSeq 500 with the read configuration as 76 bp, single end. Reads were mapped to the hg19 (UCSC version from igenomes) using Tophat and annotated using a custom R script using the UCSC known Genes table as a reference. Read counts were generated using feature Counts and the differential expression analysis was performed using edgeR (false discovery rate (FDR) < 0.05 and mRNA fold change > 1.5 as cutoffs), as described previously (17).

### Immunoblot assay

Cells were lyzed in modified lysis buffer (50 mM Tris-HCl, pH 7.5, 150 mM NaCl, 1 mM β-mercaptoethanol, 1% Igepal, and protease inhibitor cocktail) on ice for 30 min. The equal amount of proteins were separated on SDS-PAGE gel and transferred to nitrocellulose membrane. The membrane was blocked with 5% milk and incubated overnight with primary antibodies: anti-Flag antibody (Sigma, F3165), anti-MGMT antibody (Proteintech, 17195-1-AP), anti-PAR antibody (Trevigen, 4336-BPC-100), anti-PARP-1 antibody (Proteintech, 22999-1-AP), anti-γH2AX antibody (Millipore, 05-636), anti-H2AX antibody (Proteintech, 10856-1-AP), anti-phospho-Chk1 (Ser345) antibody (Cell Signaling Technology, 2348), anti-phospho-Chk1 (Ser296) antibody (Cell Signaling Technology, 2349), anti-Chk1 antibody (Santa Cruz, sc-8408), anti-phospho-Chk2 (Thr68) (C13C1) antibody (Cell Signaling Technology, 2197), anti-phospho-Chk2 (D9C6) antibody (Cell Signaling Technology, 6334), anti-PP2A antibody (Proteintech, 10321-1-AP), anti-H3K27me3 (Cell Signaling Technology, 9733), or anti-actin antibody (Proteintech, 66009-1-AP) at 4°C, followed by donkey anti-mouse or goat anti-rabbit IgG conjugated to HRP for 1 h at room temperature. After washing, the immune complexes were detected by the SuperSignal West Pico Chemiluminescent Substrate (Fisher, P134578) and imaged by ChemiDoc system (Bio-Rad).

### Immunostaining assay

Cells were fixed with 4% paraformaldehyde, permeabilized with 0.05% Triton X-100, and blocked with 3% BSA in phosphate buffered saline (PBS). Cells then were incubated overnight with anti-Flag antibody (1:2000) and anti-MGMT antibody (1:500) at 4 °C, washed with PBS with 0.1% Tween-20 for 3 times, and incubated with Alexa488 donkey anti-mouse IgG (1:1000) and Cy3 donkey anti-rabbit IgG (1:1000) for 90 min in dark. After washing for three times, cells were incubated with DAPI (1:1000) for 5 min. Cells were washed again and mounted with anti-fade mounting medium. Immunofluorescent analysis was carried out with a Zeiss Observer Z1 fluorescence microscope.

### MGMT promoter methylation assay

*MGMT* promoter methylation was determined as described previously (21). Briefly, genomic DNA was isolated from parental and KDM6B KO2 HeLa cells and subjected to bisulfite modification by EpiMark® Bisulfite Conversion Kit according to the manufacturer’s instructions (New England Biolabs, E3318S). The converted DNA was eluted in DNase/RNase-free water and examined by PCR using SYBR Green PCR Master Mix (Bio-Rad) and primers listed in Supplementary Table S2. Genomic DNA treated with a CpG Methyltransferase M.SssI (New England Biolabs) served as positive control. Genomic DNA without bisulfite conversion was used as the negative control. PCR products were separated on nondenaturated 6% polyacrylamide gels and imaged by ChemiDoc system (Bio-Rad).

### *In vitro* MGMT methyltransferase activity assay

MGMT methyltransferase activity was determined by MGMT assay kit (MD0100, Sigma-Aldrich). Briefly, scrambled control (SC) and KDM6B KO2 HeLa cells were pretreated with vehicle or BG (200 μM) for 30 min and lysed in the modified lysis buffer (50 mM Tris, pH 7.5, 1mM EDTA, 1mM DTT, 5% Glycerol, 50 mM NaCl). The resulting lysates were incubated with a 23-bp customized biotin-labeled DNA substrate containing an O^6^-methylguanine next to the PstI cleavage site (Sigma) for 2 h at 37 °C. Then the reaction was stopped at 65 °C for 5 min. The resulting DNA purified via phenol/chloroform/isoamyl alcohol (25:24:1) with 10 μg yeast tRNA serving as a carrier was incubated with PstI at 37 °C for 1 h (10 μl reaction system). The reaction was stopped by 5 μl FBXE buffer (90% Formamide, 0.1% w/v Bromophenol blue, 0.1% Xylene cyanole, 20 mM EDTA), heated at 95°C for 5 min and quickly chilled on ice, followed by separation on 20% 7 M urea SDS-PAGE gel and immunoblot assay with anti-Biotin antibody (Thermo Fisher Scientific, 692033).

### Quantitative reverse transcription-polymerase chain reaction (RT-PCR) assay

Total RNA was isolated using TRIzol reagent (ThermoFisher) and cleaned with RNase-free DNase I (ThermoFisher). cDNA was synthesized with M-MuLV Reverse Transcriptase (200 U/μL, New England Biolabs). Real-time PCR was performed by a CFX-96 real-time system (Bio-Rad) using iTaq universal SYBR green supermix (Bio-Rad) with primers listed in Supplementary Table S2. mRNA expression was quantified as described previously (17).

### Statistical analysis

Statistical evaluation was performed by unpaired two-tailed Student’s *t* test between two groups and by one- or two-way ANOVA with Tukey’s multiple comparisons, Dunnett’s multiple comparisons or Sidak’s multiple comparisons within multiple groups as indicated using GraphPad Prism 8.0 software. Data are shown as mean ± SEM. *p* < 0.05 is considered significant.

## Results

### PARP-1 is required for alkylating agent-induced cell death in cancer cells

Alkylating agents including SN1-type alkylating agent MNNG and SN2-type alkylating agent MMS induced DNA damage and cell death in cancer cells (Fig. 1; Supplementary Fig. 1A and B). To determine which cell death pathway is responsible for alkylating agent-induced cancer cell death, we screened cell death inhibitors in human cervical cancer HeLa cells treated with DMSO or MNNG (50 μM). Inhibitors for apoptosis (z-VAD, 10 μM), necroptosis (necrostatin-1, 10 μM), and autophagy (3-methyladenine [3-MA], 10 mM) all failed to inhibit MNNG-induced cell death (Fig. 1A). In contrast, MNNG-induced cell death was fully prevented by PARP inhibitor DPQ (30 μM) or Olaparib (10 μM) (Fig. 1A). PARP-1 senses DNA damage for its activation. PAR formation determined by anti-PAR antibody showed that MNNG treatment for 15 min robustly induced PARP-1 activation in HeLa cells, which was completely blocked by DPQ (30 μM) (Fig. 1B). To further confirm the role of PARP-1 in alkylating agent-induced cell death, we generated and validated PARP-1 protein knockdown (KD) in HeLa cells using its small interfering RNAs (siRNAs) (Fig. 1C). Similar to pharmacological interventions, PARP-1 KD also fully blocked MNNG-induced HeLa cell death (Fig. 1D and E).

**Figure 1.**
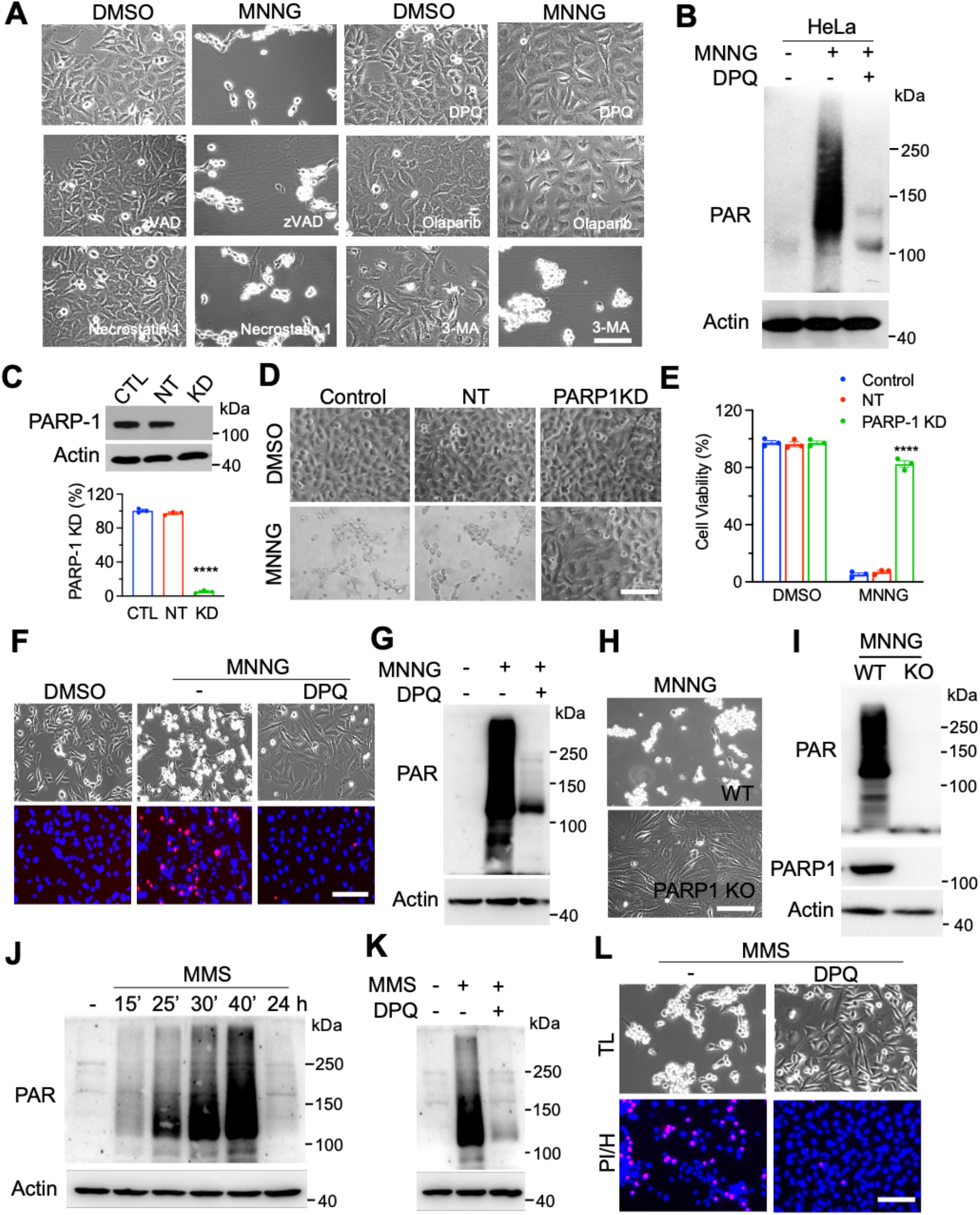
PARP-1 is required for alkylating agent-induced cancer cell death. (**A**) Representative cell death images in HeLa cells treated with DMSO or MNNG (50 μM, 15 min) for 24 h in the presence or absence of cell death inhibitors z-VAD (10 μM), necrostatin-1 (10 μM), 3-methyladenine (3-MA, 10 mM). Scale bar, 20 μm. (**B**) Immunoblot analysis of PARP-1 activation in HeLa cells treated with vehicle (−) or MNNG (50 μM) for 15 min in the presence or absence of DPQ (30 μM). (**C**) Immunoblot analysis of PARP-1 knockdown (KD) in HeLa cells (top). PARP-1 protein levels are quantified (bottom, mean ± SEM, *n* = 3). ****p < 0.0001 *vs*. control by one-way ANOVA Dunnett’s multiple comparisons test. CTL, control. NT, non-target. (**D** and **E**) Representative cell death images in control, NT, and PARP-1 KD HeLa cells treated with DMSO or MNNG for 24 h (D). Cell death is quantified in E (mean ± SEM, *n* = 3). Scale bar, 20 μm. ****p < 0.0001 *vs*. control by two-way ANOVA Sidak’s multiple comparisons test. (**F**) Representative cell death images in MDA-MB-231 cells treated with DMSO or MNNG (75 μM, 15 min) for 24 h in the presence or absence of DPQ. Scale bar, 20 μm. PI/H, propidium iodide/Hoechst staining. TL, transmission light. (**G**) Immunoblot analysis of PARP-1 activation in MDA-MB-231 cells treated with vehicle (−) or MNNG for 15 min in the presence or absence of DPQ. (**H**) Representative cell death images in wildtype (WT) and PARP-1 KO MDA-MB-231 cells treated with MNNG (75 μM, 15 min) for 24 h. Scale bar, 20 μm. (**I**) Immunoblot analysis of PARP-1 activation in WT and PARP-1 KO MDA-MB-231 cells treated with MNNG for 15 min. (**J**) Immunoblot analysis of PARP-1 activation and DNA damage in MDA-MB-231 cells treated with vehicle (−) or MMS (2 mM) for indicated time. (**K**) Immunoblot analysis of PARP-1 activation in MDA-MB-231 cells treated with vehicle (−) or MMS for 1 h in the presence or absence of DPQ. (**L**) Representative cell death images in MDA-MB-231 cells treated with MMS and/or DPQ for 24 h. Scale bar, 20 μm.

We next studied whether this PARP-1-dependent cell death is also applied to other human cancer cells. Human breast cancer MDA-MB-231 cells were treated with MNNG (100 μM) for 15 min in the presence or absence of DPQ (30 μM) and cell death was assessed by propidium iodide/Hoechst staining 24 h after treatment. In line with results in HeLa cells, MNNG treatment killed MDA-MB-231 cells, which was blocked by DPQ (Fig. 1F). Similar effects were also observed in MCF-7 and SUM159 cells (Supplementary Fig. 1C). DPQ treatment also abolished MNNG-induced PARP-1 activation in MDA-MB-231 cells (Fig. 1G). Again, genetic knockout (KO) of PARP-1 prevented MNNG-induced PAR formation and cell death in MDA-MB-231 cells (Fig. 1H and I). Finally, we studied the effect of PARP-1 on other alkylating agents-induced cell death. To this end, we treated MDA-MB-231 cells with MMS (4 mM) or vehicle. Treatment of MMS rapidly increased PAR formation in a time-dependent manner with peak observed at 40 min and caused DNA damage as γH2AX was remarkable increased 24 h after treatment (Fig. 1J, supplementary Fig. 1B). Pretreatment of DPQ (30 μM) blocked MMS-induced PARP-1 activation and cell death in MDA-MB-231 cells (Fig. 1K and L). Collectively, these findings indicate that alkylating agents activates PARP-1 to induce cancer cell death.

### Identification of KDM6B as a key cell death effector involved in PARthanatos

To unbiasedly identify key mediators controlling PARP-1-dependent cell death in response to alkylating agents, we performed a genome-wide CRISPR/Cas9 screening in HeLa cells (Fig. 2A). HeLa cells were transduced with pooled genome-wide CRISPR KO (GeCKO v2) libraries A and B containing 122,411 sgRNAs. After puromycin selection, cells were then treated with MNNG (75 μM) for 15 min to trigger PARthanatos. The survived cells resistant to PARthanatos were harvested for genomic DNA isolation and deep sequencing, which identified 18 enriched sgRNA hits (Fig. 2B). Given that two highly enriched KDM6B sgRNAs were identified from the screening, KDM6B, which is a histone H3K27 demethylase, was selected as a top candidate for further studies. To validate the screening results, we generated two independent KDM6B KO HeLa cell lines by the CRISPR/Cas9 technique and these KO cells were confirmed by Sanger sequencing (Fig. 2C). KDM6B KO1-2 significantly inhibited MNNG-induced HeLa cell death (Supplementary Fig. 2A and B).

**Figure 2.**
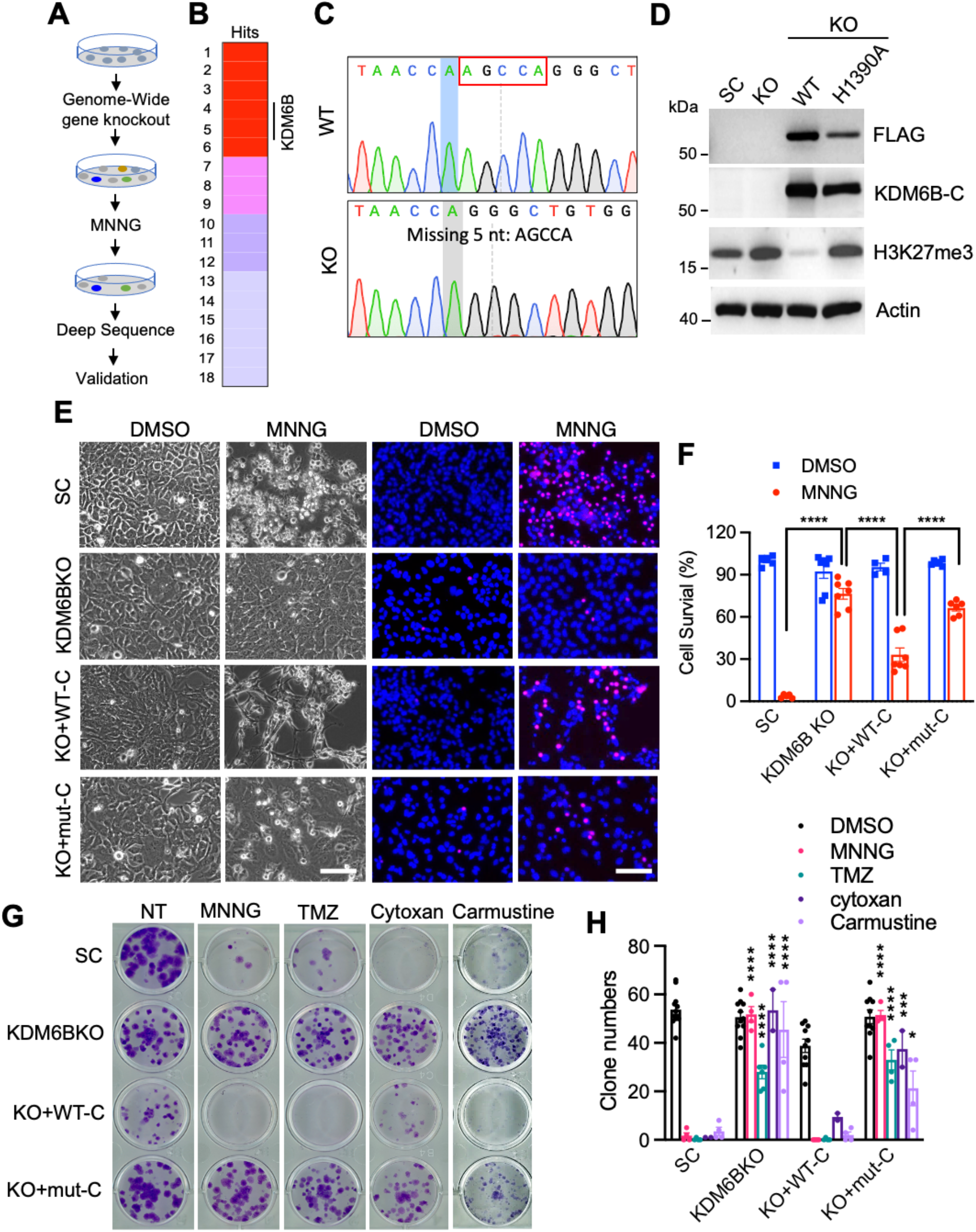
Identification of KDM6B as a key regulator of PARP-1-dependent cell death. (**A**) The scheme of the CRISPR screening. (**B**) Top 18 hits including KDM6B were identified from the screening. Red, > 105,000 reads. Pink, > 200 reads. Purple, >10 reads. Gray, ≥ 1 read. (**C**) Genotyping of KDM6B KO HeLa cells. (**D**) Immunoblot analysis of KDM6B KO and rescued HeLa cells. (**E** and **F**) Representative cell death images in SC, KDM6B KO2, and rescued HeLa cells 24 h after the treatment with DMSO or MNNG (50 μM, 15 min). (E). PI-positive cells are quantified in F (mean ± SEM, *n* = 4-7). Scale bar, 20 μm. PI/H, propidium iodide/Hoechst staining. TL, transmission light. ****p < 0.0001 by two-way ANOVA Sidak’s multiple comparisons test. (**G** and **H**) Representative colony survival in SC, KDM6B KO2, and rescued HeLa cells treated with vehicle, MNNG (2 μM), TMZ (250 μM), Cytoxan (500 μM), and Carmustine (25 μM) for 10 days (G). Colony numbers are quantified in H (mean ± SEM, *n* = 2-12). *p < 0.05; ***p < 0.001; ****p < 0.0001 *vs*. DMSO by two-way ANOVA Tukey’s multiple comparisons test.

To determine whether the demethylase activity of KDM6B is required for alkylating agent-induced cell death, we next generated KDM6B rescue cells expressing wildtype (WT) or catalytically inactive (H1390A) C-terminal KDM6B truncate (KDM6B-C, Fig. 2D)(22). KDM6B KO2 increased H3K27me3 in HeLa cells (Supplementary Fig. 2C). In line with previous studies (23), WT but not H1390A KDM6B-C showed the demethylase activity because it eliminated the level of H3K27me3 in HeLa cells (Fig. 2D), indicating the specificity of these KO and rescued cell lines. As expected, MNNG treatment robustly induced HeLa cell death, which was blocked by KDM6B KO2 (Fig. 2E and F). Ectopic expression of WT KDM6B-C sensitized KO2 cells to MNNG treatment, whereas catalytically inactive KDM6B-C mutant failed to do so (Fig. 2E and F).

Clonogenic assay showed that KDM6B KO preserved HeLa cell survival against MNNG treatment, which was reversed by WT but not mutant KDM6B-C (Fig. 2G and H). Similarly, alkylating agents, which are currently used in the clinic, including TMZ (250 μM), Cytoxan (500 μM), and Carmustine (25 μM), significantly reduced colony numbers. KDM6B KO conferred HeLa cells resistance to the treatment of these alkylating agents (Fig. 2G and H). Overexpression of WT, but not catalytically inactive KDM6B-C H1390A mutant, re-sensitized HeLa cells to treatment of these alkylating agents (Fig. 2G and H). In parallel, we also generated HeLa cells expressing empty vector (EV), full-length WT KDM6B or catalytically inactive mutant (Supplementary Fig. 2D and E). Consistent with KDM6B-C H1390A, full-length KDM6B H1390A mutant lost its demethylase activity as indicated by high H3K27me3 staining in cells and blocked MNNG-induced cell death in HeLa cells (Supplementary Fig. 2E and F). Taken together, these findings identified KDM6B as a key cell death effector in PARthanatos, which requires its demethylase activity.

### KDM6B suppresses MGMT expression via its demethylase activity and regulates alkylating agent-induced PARP-1 activation and DNA damage

To dissect the mechanism by which KDM6B facilitates PARP-1-dependent cell death in response to alkylating agents, we analyzed KDM6B transcriptome in parental, KDM6B KO2, and rescued WT or mutant KDM6B HeLa cells by RNA-sequencing. KDM6B upregulated 811 genes but downregulated 690 genes (FDR < 0.05 and fold change > 2 as cutoffs, Fig. 3A). 290 out of 811 upregulated genes and 250 out of 690 downregulated genes were rescued by WT KDM6B (fold change > 1.5 as cutoff, Fig. 3B). Importantly, the majority of rescued genes were regulated by KDM6B demethylase activity (Fig. 3C). In parallel with RNA-sequencing study, we found that KDM6B regulated PARP-1 activation following MNNG treatment in a time-dependent manner in HeLa SC cells (Fig. 3D). KDM6B KO significantly reduced PARP-1 activation and PAR formation (Fig. 3D). In line with that, DNA damage marker γH2AX was also elevated in a time-dependent manner after MNNG treatment, which was greatly inhibited in KDM6B KO2 cells (Fig. 3D). MGMT is a key de-alkylating enzyme that transfers methyl groups from the O^6^ position of guanine to its own active residue cysteine 145 leading to a direct repair of guanine, and subsequently methylated MGMT protein is degraded in the proteasome (24). Thus, MGMT was identified as the top hit among KDM6B demethylase activity-regulated genes, which is likely to regulating DNA damage response, PARP-1 activation and the resistance of KDM6B KO cells to alkylating agents (Fig. 2E-H, supplementary Fig. 2A, B, F).

**Figure 3.**
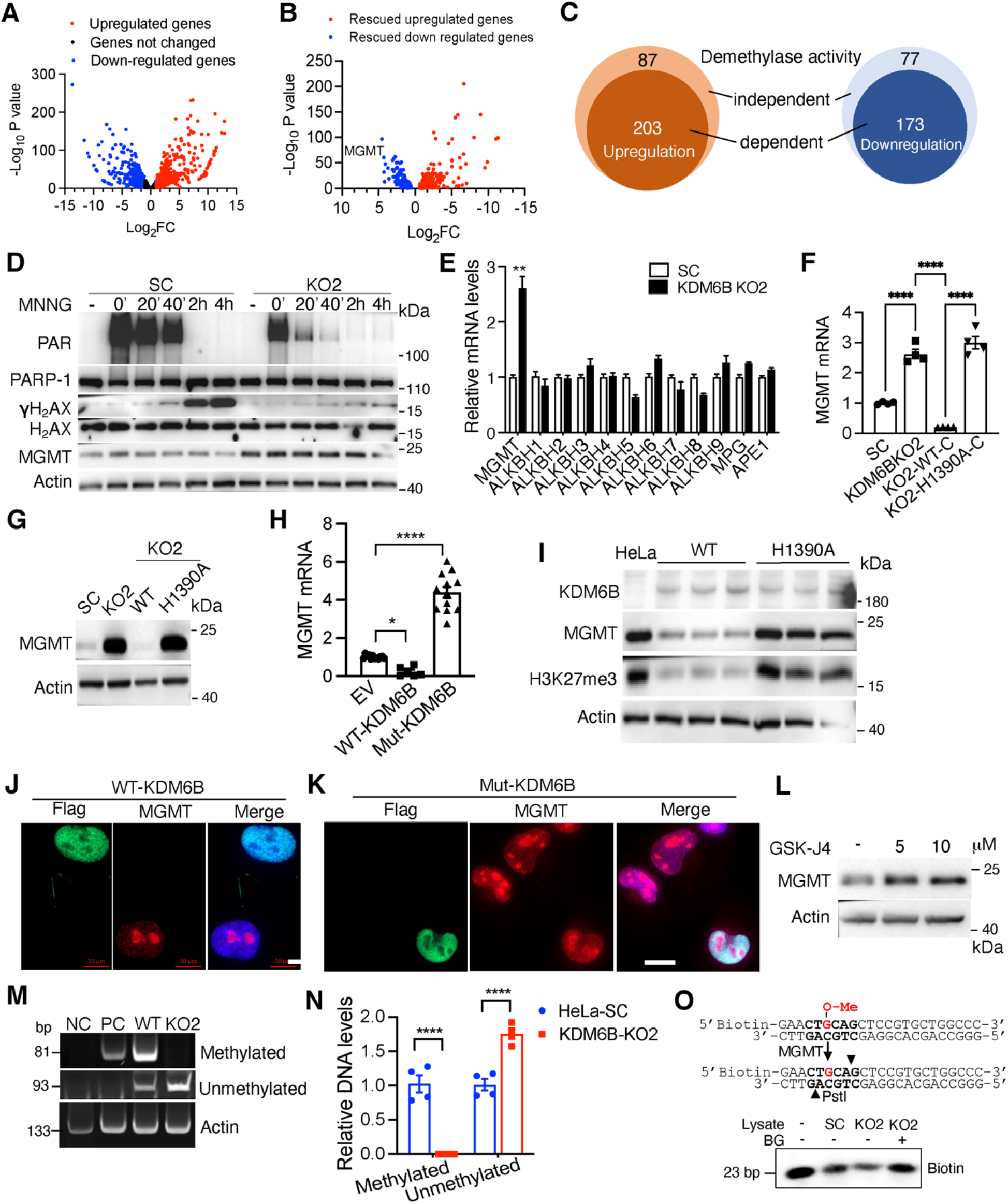
KDM6B regulates PARP-1 activation and MGMT expression. (**A** and **B**) Volcano plots of KDM6B target genes in HeLa cells. *n* = 2. (**C**) Venn diagram of KDM6B rescued target genes in HeLa cells. (**D**) Immunoblot analysis of indicated proteins in SC and KDM6B KO2 HeLa cells treated with vehicle (−) or MNNG for indicated time. (**E**) RT-qPCR analysis of indicated genes in SC and KDM6B KO2 HeLa cells (mean ± SEM, *n* = 3-6). **p < 0.01 by unpaired two-tailed Student’s *t* test. (**F**) RNA-seq analysis of MGMT expression in SC, KDM6B KO2, and rescued HeLa cells (mean ± SEM, *n* = 4). ****p < 0.0001 by one-way ANOVA Sidak’s multiple comparisons test. (**G**) Immunoblot analysis of MGMT in SC, KDM6B KO2, and rescued HeLa cells. (**H**) RT-qPCR analysis of MGMT expression in HeLa cells expressing EV, WT KDM6B, or catalytically mutant (mut) KDM6B (mean ± SEM, *n* = 6-12). *p < 0.05; ****p < 0.0001 *vs*. EV by one-way ANOVA Dunnett’s multiple comparisons test. (**I**) Immunoblot analysis of indicated proteins in HeLa cells expressing WT or H1390A KDM6B. (**J** and **K**) Representative KDM6B and MGMT immunostaining images in HeLa cells expressing WT or catalytically mutant KDM6B. Scale bar, 20 μm. (**L**) Immunoblot analysis of MGMT in A549 cells treated with or without GSK-J4 for 24 h. (**M** and **N**) Representative agarose gels of methylated and unmethylated MGMT promoters in WT and KDM6B KO2 HeLa cells (M). DNA intensity is quantified in N (mean ± SEM, *n* = 4). ****p < 0.0001 by two-way ANOVA Sidak’s multiple comparisons test. (**O**) *In vitro* MGMT activity assay. The assay strategy is shown on the top. Immunoblot analysis of biotin is shown at the bottom.

To validate RNA-seq results, we performed RT-qPCR assay in SC and KDM6B KO2 HeLa cells and found KDM6B KO2 remarkably increased MGMT mRNA expression, which was not altered upon MNNG treatment (Fig. 3E; Supplementary Fig. 3A). MGMT protein levels were also upregulated in KDM6B KO2 HeLa cells but declined 4 h after MNNG treatment (Fig. 3D), which is consistent with the previous finding that MGMT is degraded at the proteasome after repairing damaged DNA. In contrast, other nucleic acid demethylases ALKBH1-9, DNA glycosylase MPG, and endonuclease APE1, which are also known to be involved in alkylating agent-induced DNA damage (16), were not regulated by KDM6B in HeLa cells (Fig. 3E).

Further, our rescue study showed that expression of WT KDM6B-C but not H1390A mutant reversed induction of MGMT mRNA and protein levels conferred by KDM6B KO2 in HeLa cells (Fig. 3F and G). Similarly, ectopic expression of full-length KDM6B but not H1390A mutant inhibited MGMT mRNA and protein expression (Fig. 3H and I). Immunostaining assay further confirmed inhibition of MGMT expression at the nucleus by KDM6B through its demethylase activity (Fig. 3J and K). We next employed KDM6B inhibitor GSK-J4 and found that treatment of GSK-J4 (5 or 10 μM) robustly increased MGMT protein levels in A549 cells (Fig. 3L). In line with these *in vitro* results, KDM6B expression was also negatively correlated with MGMT expression in human breast and brain tumors (Supplementary Fig. 3B and C). These findings support that KDM6B suppresses MGMT expression via its demethylase activity.

### KDM6B regulates DNA methylation at the *MGMT* promoter and inhibits MGMT methyltransferase activity

To understand the mechanism underlying KDM6B-repressed *MGMT* transcription, we assessed whether KDM6B controls DNA methylation at the *MGMT* promoter. Strong DNA methylation was detected at the *MGMT* promoter in WT HeLa cells, but not KDM6B KO2 cells (Fig. 3M and N). Consistently, the PCR product of the unmethylated *MGMT* promoter was significantly increased in KDM6B KO2 cells (Fig. 3M and N). Together, these findings indicate that KDM6B increases methylation of *MGMT* promoter to repress MGMT expression.

We next studied whether KDM6B inhibits the methyltransferase activity of MGMT. Whole cell lysates from KDM6B KO2 and SC HeLa cells were incubated with a biotin-labeled DNA substrate with a methylated guanine at its O^6^-position. Once the methylated DNA substrate is converted by MGMT into un-methylated form, it can be cut by the restriction enzyme PstI, leading to reduced biotin signal. We found that biotin-labeled DNA levels were decreased when incubated with KDM6B KO2 cell lysates as compared with SC cell lysates, which was completely reversed by a specific MGMT inhibitor O^6^-benzylguanine (BG), which is a pseudo substrate for MGMT (Fig. 3O). These results indicate that KDM6B KO cells have a higher MGMT methyltransferase activity than SC cells, which may be due to increased MGMT expression.

### MGMT inhibition re-sensitizes KDM6B KO cells to alkylating agents through PARP-1 activation

To determine the role of MGMT in KDM6B KO-induced resistance to PARthanatos in response to alkylating agents, we treated SC and KDM6B KO2 HeLa cells with MGMT inhibitor BG (200 μM) with or without MNNG. As expected, treatment of BG for 24 h specifically depleted MGMT protein in HeLa cells, as MNNG treatment did (Fig. 4A). MNNG effectively caused death of SC but not KDM6B KO2 HeLa cells 24 h after treatment (Fig. 4B and C). MGMT inhibition by BG treatment robustly killed MNNG-resistant KDM6B KO2 HeLa cells at 72 h, but not 24 h, after MNNG treatment (Fig. 4B and C). Treatment of BG alone for up to 96 h had the marginal effect on death of both SC and KDM6B KO2 cells (Fig. 4B and C). In line with pharmacological inhibition, genetic deletion of MGMT also significantly reversed HeLa cell survival conferred by KDM6B loss under MNNG treatment conditions (Fig. 4D-F). In contrast, MNNG-induced cell death was significantly reduced by overexpression of WT MGMT but not DNA methyltransferase inactive MGMT C145A mutant (Supplementary Fig. 4A and B). In parallel, we found that BG treatment reduced endogenous MGMT protein levels and re-sensitized KDM6B KO2 cells to MMS (Supplementary Fig. 4C and D). However, MMS treatment alone did not affect MGMT or MPG protein levels (Supplementary Fig. 4E). Together, these findings indicate that MGMT contributes to KDM6B KO cells resistance to alkylating agents.

**Figure 4.**
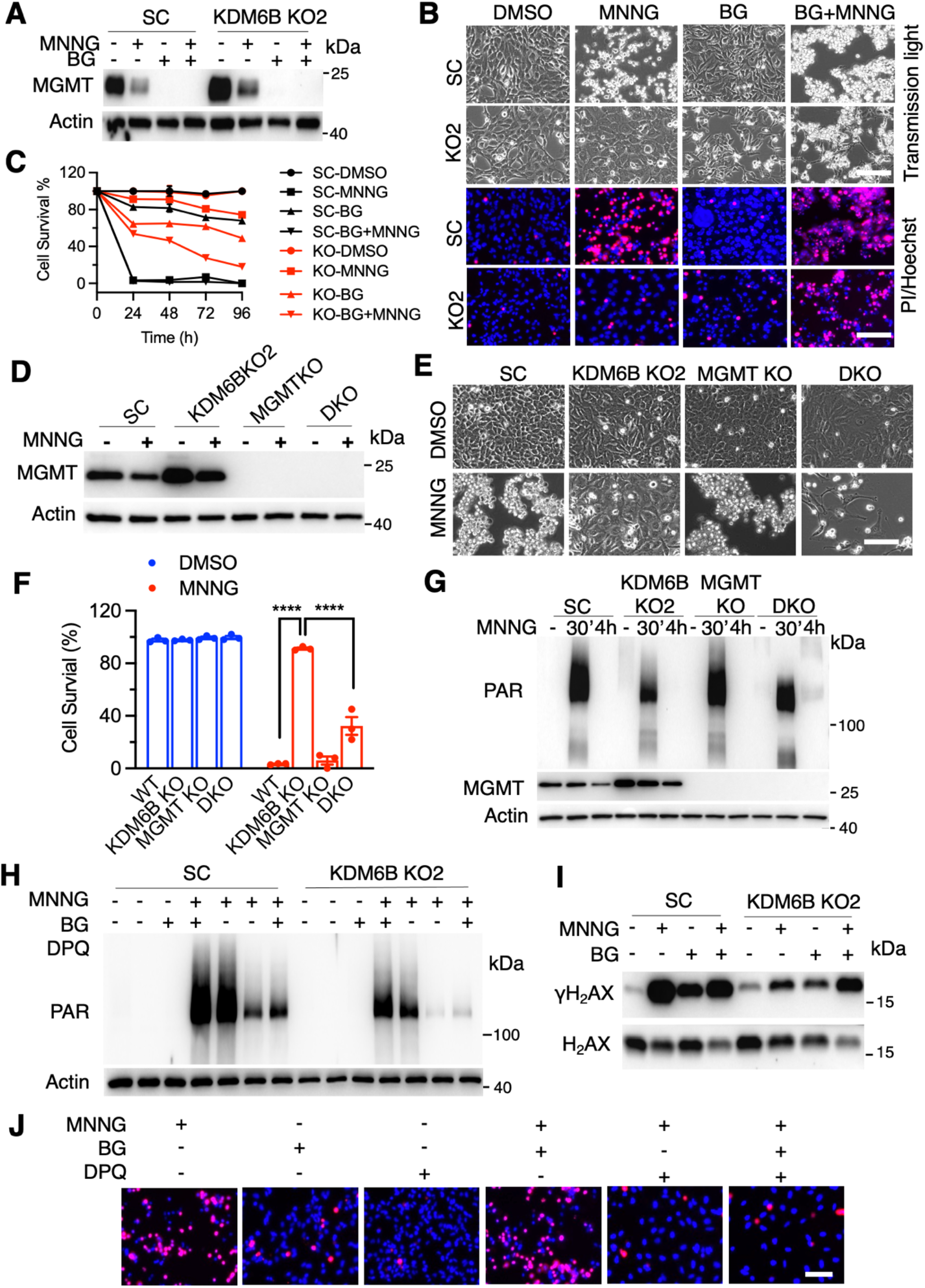
Pharmacological inhibition of MGMT or deletion of MGMT sensitizes KDM6B KO cells to alkylating agents. (**A**) Immunoblot analysis of MGMT in SC and KDM6B KO2 HeLa cells 24 h after the treatment of MNNG (50 μM, 15 min) and/or BG (200 μM). (**B** and **C**) Representative cell death images in SC and KDM6B KO2 HeLa cells treated with MNNG and/or BG (B) and PI-positive cells are quantified in C (mean ± SEM, *n* = 3). (**D**) Immunoblot analysis of MGMT in SC, KDM6B KO2, MGMT KO, and KDM6B/MGMT DKO HeLa cells 24 h after MNNG treatment. (**E** and **F**) Representative cell images in SC, KDM6B KO2, MGMT KO, KDM6B/MGMT DKO HeLa cells 24 h after MNNG treatment (E). Cell death is quantified in F (mean ± SEM, *n* = 3). ****p < 0.0001 by two-way ANOVA Sidak’s multiple comparisons test. (**G**) immunoblot analysis of PARP-1 activation in SC, KDM6B KO2, MGMT KO, KDM6B/MGMT DKO HeLa cells treated with vehicle (−) or MNNG for indicated time. (**H**) Immunoblot analysis of PARP-1 activation in SC and KDM6B KO2 HeLa cells treated with MNNG, BG, and/or DPQ for 15 min. (**I**) Immunoblot analysis of DNA damage in SC and KDM6B KO2 HeLa cells 24 h after MNNG and/or BG treatment. (**J**) Representative cell death images in HeLa cells 24 h after treatment of MNNG, BG, and/or DPQ.

We next studied whether MGMT repairs DNA to block PARP-1 activation, leading to KDM6B KO cells resistance to alkylating agents. MNNG increased PAR levels in SC cells and PAR levels were robustly decreased in KDM6B KO2 HeLa cells 30 min after MNNG treatment, which was partially rescued in KDM6B and MGMT double KO (DKO) cells (Fig. 4G). Weak PAR was even detected in DKO cells 4 h after MNNG treatment (Fig. 4G). MGMT KO caused comparable levels of PAR formation as that of SC in response to MNNG (Fig. 4G). Consistently, PAR reduction in KDM6B KO2 HeLa cells treated with MNNG was also partially reversed by BG treatment, which was almost completely eliminated by DPQ (Fig. 4H). In line with PARP-1 activation, MNNG treatment increased DNA damage in SC cells indicated by γH2AX levels and KDM6B KO reduced γH2AX levels (Fig. 4G), which was reversed by MGMT inhibition induced by BG treatment (Fig. 4I). Finally, we found that treatment of DPQ prevented cell death from MNNG or MNNG plus BG in HeLa cells (Fig. 4J). These results indicate that KDM6B/MGMT are the upstream regulators of PARP-1 controlling DNA damage and alkylating agent-induced cell death.

### KDM6B suppresses alkylating agent-induced prolonged activation of Chk1 kinase and its associated DNA repair

Our results above showed that death of KDM6B KO cells upon BG treatment was significantly delayed as compared with that of SC cells and KDM6B KO cells were also resistant to MMS treatment that is not expected to increase O6-methylguanine levels. These findings suggest that KDM6B KO might trigger a second mechanism independent of MGMT conferring cells resistance to alkylating agents-induced cell death, especially when DNA alkylation evades from MGMT-mediated direct DNA repair. To determine whether DNA damage checkpoint response is involved in resistance to alkylating agents in KDM6B KO cells, we studied if KDM6B regulates phosphorylation of Chk1, which regulates DNA damage response and cell cycle checkpoint response, after exposure to MNNG. We found that Chk1 was phosphorylated at both Ser345 and Ser296 upon MNNG treatment but quickly declined 4 h after treatment in SC cells (Fig. 5A). In contrast, constitutive phosphorylation of Chk1 up to at least 6 h was observed in KDM6B KO2 HeLa cells after MNNG treatment (Fig. 5A). Chk1 activation has been known to phosphorylate its downstream factor Cdc25A, a step required for Cdc25A ubiquitination and degradation, thereby leading to cell cycle arrest (25). In line with acute Chk1 activation in SC cells, Cdc25A protein was quickly degraded while DNA damage was quickly increased as indicated by γH2AX in SC cells from 1 h to 4 h after MNNG treatment, indicating cell cycle arrest and deficiency of DNA repair (Fig. 5A). In contrast, levels of Cdc25A remained relatively stable in KDM6B KO2 HeLa cells at the late period (2-6 h after MNNG treatment) and DNA damage was not further accumulated from 1 h to 4 h after MNNG treatment, indicating that prolonged Chk1 activation enhanced checkpoint associated repair without causing cell cycle arrest (Fig. 5A). Distinct to phospho-Chk1, Chk2 phosphorylation showed a similar pattern in both SC and KDM6B KO2 HeLa cells (Fig. 5A). The expression of protein phosphatase 2A (PP2A), which is involved in Cdc25A dephosphorylation, was not obviously altered in KDM6B KO2 HeLa cells after MNNG treatment (Fig. 5A), excluding a possibility of increased dephosphorylation of Cdc25A at the late period in KDM6B KO cells. These data indicate that KDM6B KO causes prolonged phosphorylation of Chk1 and enhances DNA repair.

**Figure 5.**
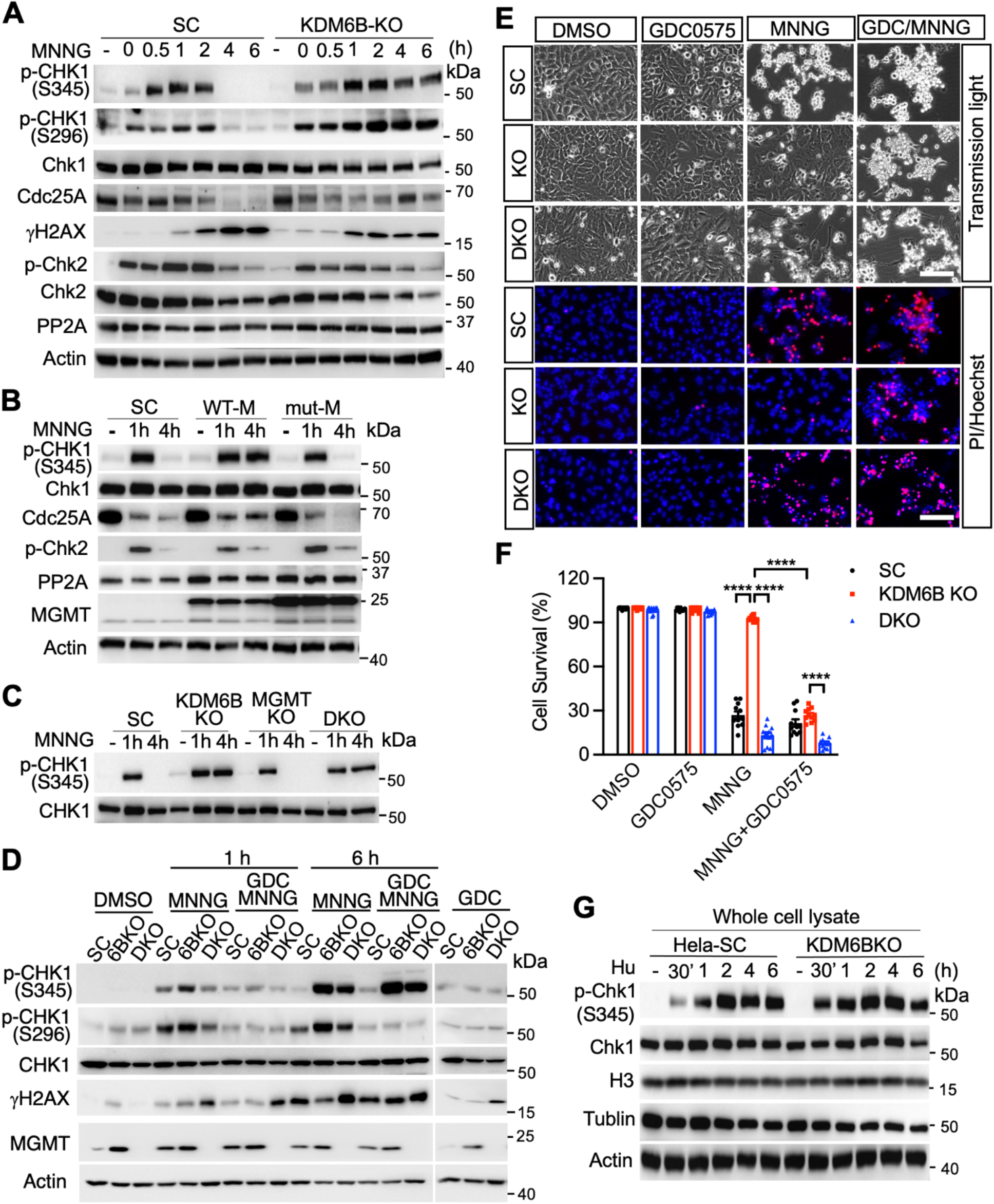
KDM6B KO promotes sustained Chk1 activation for DNA repair following alkylating agent treatment. (**A**) Immunoblot analysis of checkpoint response and DNA damage in SC and KDM6B KO2 HeLa cells 0-6 h after MNNG treatment. (**B** and **C**) Effects of MGMT and MGMT C145A mutant overexpression (B) and MGMT KO (C) on checkpoint response in SC and KDM6B KO HeLa cells 1 h and 6 h after MNNG treatment. (**D**) Effects of checkpoint inhibition by GDC0575 (50 nM) on checkpoint response and DNA damage in SC, KDM6B KO2, and KDM6B/MGMT DKO HeLa cells 1 h and 6 h after MNNG treatment. (**E** and **F**) Effects of checkpoint inhibition by GDC0575 (50 nM) on MNNG-induced cell death in SC, KDM6B KO2, and KDM6B/MGMT DKO HeLa cells 24 h after MNNG treatment. Representative cell death images are in SC, KDM6B KO2, KDM6B/MGMT DKO HeLa cells 24 h after MNNG treatment (E). Cell death is quantified in F (mean ± SEM, *n* = 3). ****p < 0.0001 by two-way ANOVA Tukey’s multiple comparisons test. (**G**) Immunoblot analysis of checkpoint response in SC and KDM6B KO2 HeLa cells 0-6 h after Hu (2 mM) treatment.

We next assessed whether MGMT alters MNNG-induced Chk1 phosphorylation by overexpression of WT-MGMT or its methyltransferase inactive C145A mutant (26). Overexpression of WT-MGMT mimicked the effects of KDM6B KO and was sufficient to cause prolonged activation of Chk1, whereas mutant MGMT C145A failed to do so (Fig. 5B). In line with prolonged Chk1 activation in WT-MGMT expressing cells, the level of Cdc25A at 4 h after MNNG treatment was similar to that at 2 h after MNNG treatment. However, Cdc25A levels were reduced in SC and MGMT C145A mutant cells at 4 h after MNNG treatment (Fig. 5B). To further study whether MGMT is necessary for the prolonged activation of Chk1, we knocked out MGMT in KDM6B KO cells and found that MGMT KO failed to reverse KDM6B KO-induced prolonged activation of Chk1 (Fig. 5C), indicating that other factors other than MGMT might contribute to DNA repair and prevent cell death in response to alkylating agents.

To study the role of checkpoint associated DNA repair besides MGMT-mediated direct DNA repair in KDM6B KO-induced resistance to alkylating agents, DNA damage was determined in KDM6B KO and KDM6B/MGMT DKO cells in the presence or absence of the specific Chk1 inhibitor GDC0575 (27) that blocks Chk1 activation thereby bypassing the checkpoint response and its associated DNA repair. As expected, KDM6B KO cells showed reduced DNA damage and cell death induced by MNNG treatment (Fig. 5D-F). MGMT KO clearly increased γH2Ax levels 1-6 h after MNNG treatment and elevated subsequent cell death (Fig. 5D-F), indicating that MGMT indeed plays a role in the resistance of KDM6B KO to alkylating agents. GDC0575 treatment inhibited Chk1 activation via reducing the phosphorylation of Chk1 at Ser296 position in SC, KDM6B KO and KDM6B/MGMT DKO cells at both 1 h and 6 h after MNNG treatment, although GDC0575 treatment did not obviously reduced the phosphorylation of Chk1 at Ser345 position, which is usually phosphorylated by its upstream factor ataxia-telangiectasia-mutated-and-Rad3-related kinase (Fig. 5D). The treatment of GDC0575 clearly increased DNA damage in KDM6B KO cells (Fig. 5D). In line with this, inhibition of Chk1 activation by GDC0575 also significantly increased the sensitivity of KDM6B KO cells to MNNG-induced cell death (Fig. 5E, F), indicating that the sustained Chk1 kinase activity-associated DNA repair also mediates KDM6B KO cell resistance to alkylating agents. Interestingly, the effects of KDM6B were specific to alkylating agents as both KDM6B WT and KO cells showed no difference of Chk1 phosphorylation in response to hydroxyurea (HU) treatment (Fig. 5G).

Taken together, KDM6B specifically suppresses alkylating agents-induced prolonged Chk1 activation and its associated DNA repair thereby promoting PARthanatic cell death in response to alkylating agents.

## Discussion

In the present study, we dissected the PARP-1-dependent cell death pathway in response to alkylating agents and identified the epigenetic regulator KDM6B as a key cell death effector required for alkylating agent-induced cell death in cancer cells. PARthanatos is one type of programmed necrotic cell death distinct from apoptosis, necrosis, necroptosis and autophagy. PARP-1 is activated when it detects DNA damage and plays a central role in PARthanatic cell death pathway. Genetic deletion or pharmacological inhibition of PARP-1 blocks alkylating agent-induced PARthanatos, which cannot be prevented by inhibitors for apoptosis, necrosis, necroptosis and autophagy.

KDM6B is a well-known histone demethylase that demethylates the gene repressive marker H3K27me3 (28). We found here that KDM6B is a key upstream regulator of PARP-1 and controls PARP-1 activation in response to alkylating agents. KDM6B suppresses MGMT expression and activity, which is required for the direct removal of DNA adduct upon alkylating agent treatment, leading to accumulation of DNA damage in cancer cells. Importantly, we showed that KDM6B increases *MGMT* promoter methylation and blocks MGMT transcription through its demethylase activity. Our results suggest that KDM6B may increase the chromatin accessibility at the *MGMT* promoter to facilitate the binding of DNA methyltransferase and subsequent methylation of the *MGMT* promoter in cancer cells. In line with our findings, a recent report showed lack of *MGMT* promoter methylation in H3.3K27M mutant glioma, in which H3K27me3 enrichment is increased locally at hundreds of genes (29). Alternatively, KDM6B may demethylate and increase the level of DNA methyltransferases, which methylates and suppresses MGMT expression. Nevertheless, our rescue as well as inactive demethylase studies showed that KDM6B demethylase activity is critical for MGMT suppression, which has been further supported by the negative correlation of their expression in human breast and brain cancers.

MGMT contributes to the first step in DNA repair by direct removal of DNA adducts added by alkylating agents. Inhibition of MGMT with BG treatment and genetic deletion of MGMT both increases DNA damage and PARP-1 activation and re-sensitizes cancer cells to alkylating agent-induced cell death, supporting the importance of MGMT in KDM6B KO cell resistance to alkylating agents. Interestingly, we found that death of KDM6B KO cells with BG treatment is significantly delayed as compared with that of SC cells, suggesting that, besides MGMT, a second independent DNA repair mechanism might also contribute to KDM6B KO cell resistance to alkylating agents, when DNA alkylation evades from MGMT-mediated direct DNA repair. Indeed, the treatment of alkylating agents in KDM6B KO cells induces sustained Chk1 activation and checkpoint response associated DNA repair thereby preventing DNA damage accumulation and further cell cycle arrest-caused cell death. In contrast, the treatment of alkylating agents in SC cells causes transient acute Chk1 phosphorylation, DNA damage increase and eventually cell cycle arrest as indicated by levels of Cdc25A and γH2Ax. These data support that enhanced checkpoint response-associated DNA repair is a 2^nd^ step in DNA repair to confer KDM6B KO cell resistance to alkylating agents. Interestingly, overexpression of MGMT promotes the prolonged activation of Chk1 mimicking the effect of KDM6B KO, whereas MGMT KO cannot reverse the sustained activation of Chk1 induced by KDM6B KO, indicating that MGMT is sufficient but not necessary to activate checkpoint associated DNA repair contributing to KDM6B KO resistance to alkylating agents.

Accumulating evidence supports that PARP-1 facilitates DNA repair in response to mild DNA damage. Blockage of PARP-1 activity sensitizes cancer cells to death. This concept has been well supported by PARP inhibitors used in the clinic for cancer therapy (30). In contrast, in response to severe DNA damage, excessive activation of PARP-1 causes large DNA fragments and caspase-independent cell death designated PARthanatos, which occurs in many organ systems and is widely involved in different neurologic and non-neurologic diseases, including ischemia-reperfusion injury after stroke and myocardial infarction, glutamate excitotoxicity, neurodegenerative diseases, inflammatory injury, reactive oxygen species–induced injury (7,31). This type of cell death is profoundly prevented by pharmacological inhibition or genetic deletion of PARP-1. The importance of PARP-1 in neuronal cell death and cancer cell death has also been strongly supported by our previous as well as current studies (7–9,31). Our current study identified the epigenetic factor KDM6B as a key cell death effector of PARthanatos and a key regulator to switch PARP-1 function and signaling from DNA repair/cell survival to DNA damage/cell death addressing a huge knowledge gap of PARP-1 functions under different contexts.

Alkylating agents are the most common chemotherapy drugs currently used in the clinic to kill cancer (10–12). However, cancer cell resistance to alkylating agents remains to be one of the biggest challenges in the clinic. We showed that KDM6B deficiency enhances MGMT-mediated DNA repair and checkpoint associated DNA repair (Figs. 3–5), thereby blocking DNA damage and PARP-1 hyperactivation, which confers cancer cell resistance to alkylating agents. Our study provides evidence of KDM6B functions in regulation of DNA repair, PARP-1 activation and cell death sensitivity to alkylating agents.

Collectively, our study elucidates an epigenetic mechanism underlying PARP-1 activation and cell death in response to alkylating agents (Fig. 6). KDM6B is a key cell death effector of PARthanatos via regulating two steps of sequential DNA repair including MGMT-mediated direct DNA repair and checkpoint associated DNA repair (Fig. 6). Low levels of KDM6B first increase direct DNA repair via elevating MGMT expression and further promote sustained checkpoint activation and its associated DNA repair, both of which confer cell resistance to alkylating agents. In contrast, high levels of KDM6B suppress MGMT-mediated DNA repair and checkpoint associated DNA repair, thereby causing PARP-1 hyperactivation and cell death. KDM6B may serve as a biomarker to predict the sensitivity of cancer cells to alkylating agents. Targeting both MGMT and checkpoint associated DNA repair might be a potential therapeutic strategy to re-sensitize cancer cells to alkylating agent treatment.

**Figure 6.**
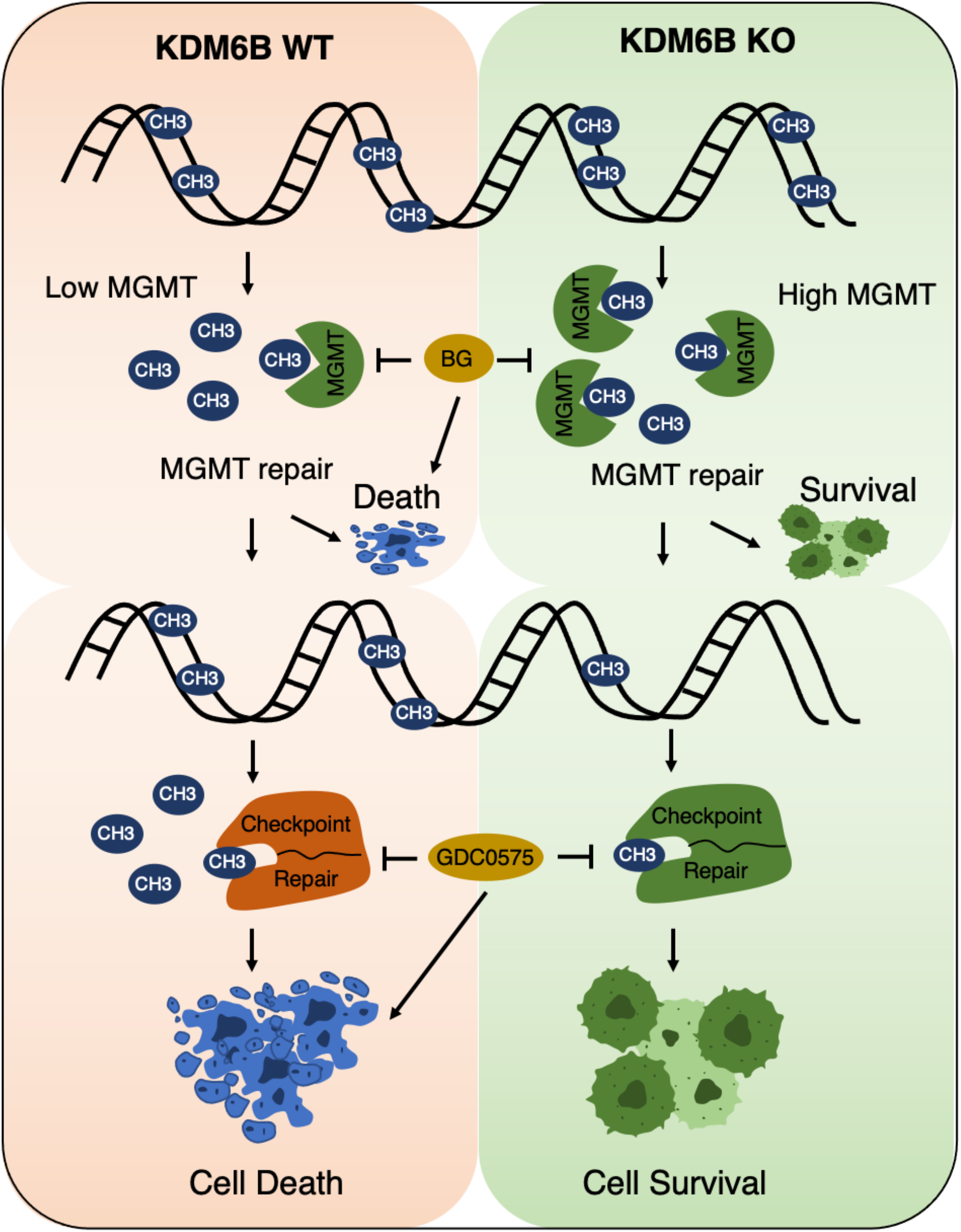
KDM6B is a key cell death effector in PARthanatos induced by alkylating agents. KDM6B regulates two sequential DNA repair processes including MGMT direct DNA repair and checkpoint associated repair. Loss of KDM6B on one hand enhances MGMT direct DNA repair to promote cell survival, on the other hand triggers sustained Chk1 activation and increases checkpoint associated DNA repair to fix DNA evading from the first step MGMT repair process leading to cell survival. High levels of KDM6B suppress both DNA repair processes leading to cell death. Blocking MGMT and checkpoint associated repair re-sensitizes cancer cells to alkylating agents.

## Declarations

### Ethics approval and consent to participate

Not applicable

### Consent for publication

Not applicable

### Availability of supporting data

The datasets used and/or analyzed during the current study are included in this article.

### Authors’ contributions

Y.W. and W. L. conceived the idea, designed research, analyzed the data and wrote the paper; M.Y. and C.W. performed most experiments. M.Z. performed RNA-seq. L.B. performed initial cell death assays and CRISPR screening validation. Y.N.W. constructed KDM6B overexpression vectors. A.K. and X.C. performed CRISPR screening and RNA-seq bioinformatics analysis. All authors read and approved the final manuscript.

## Supplementary data

**Table 1.**
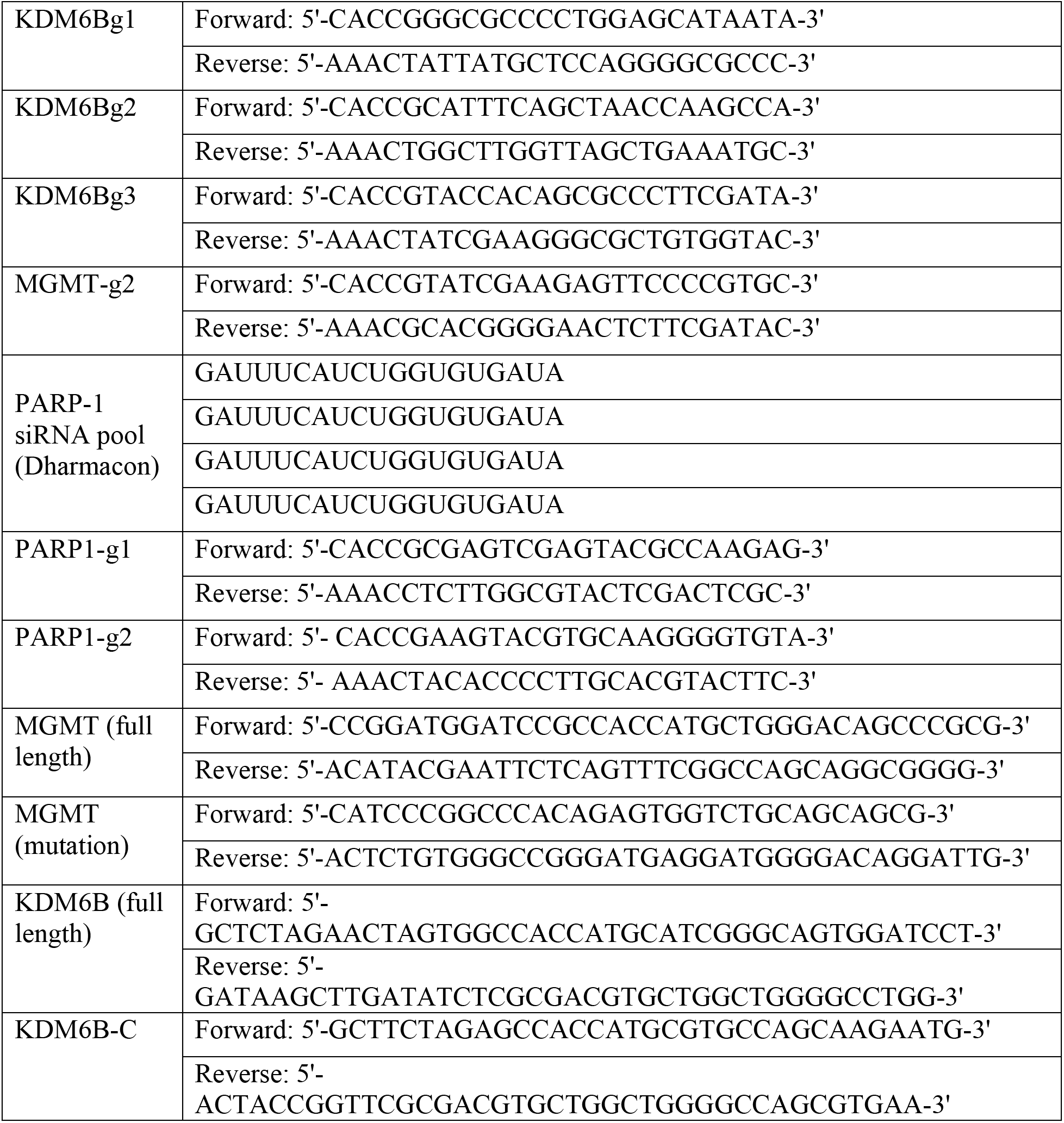
Sequences of sgRNAs, siRNAs, and primers for plasmid construction.

**Table 2.**
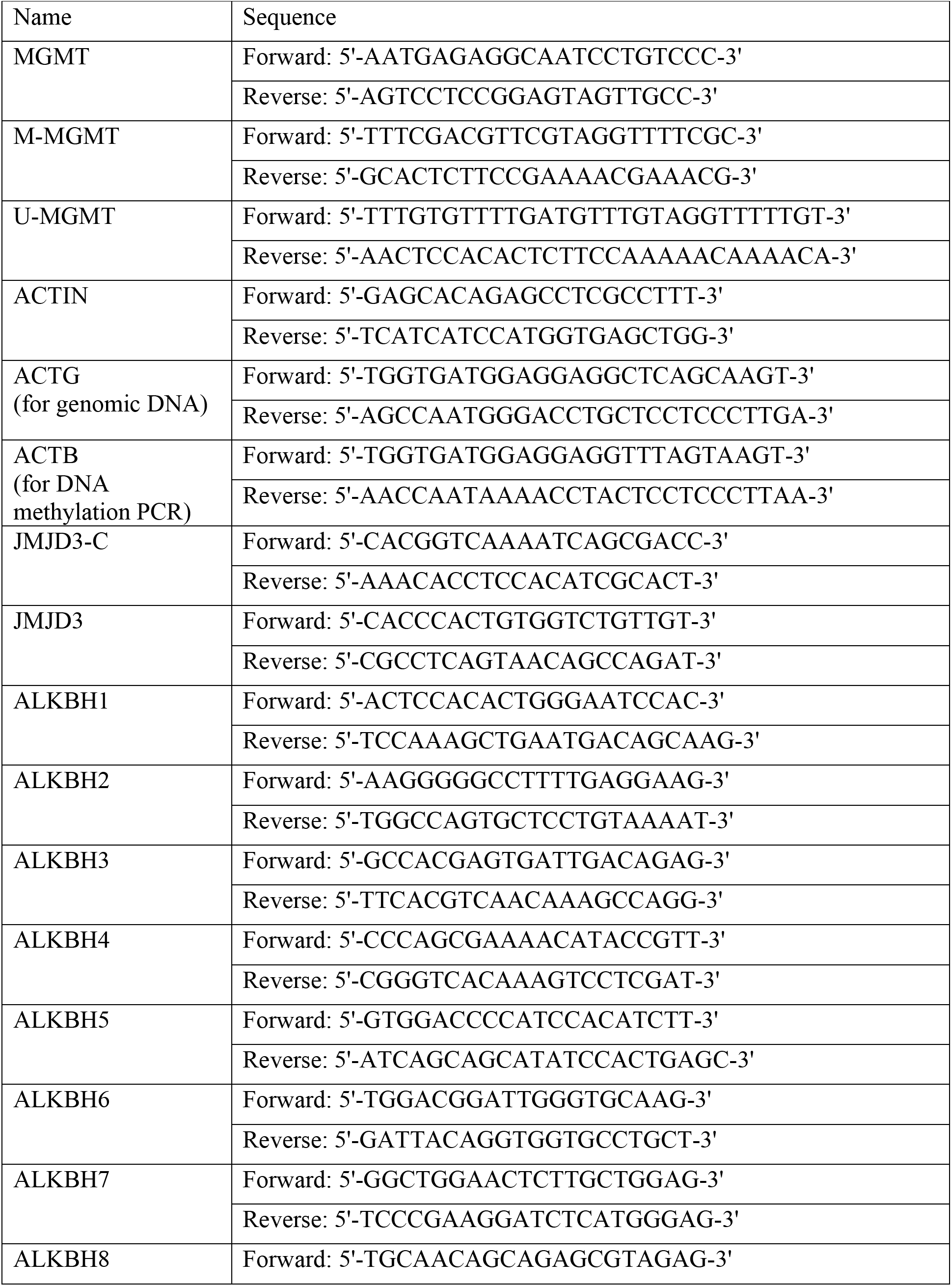
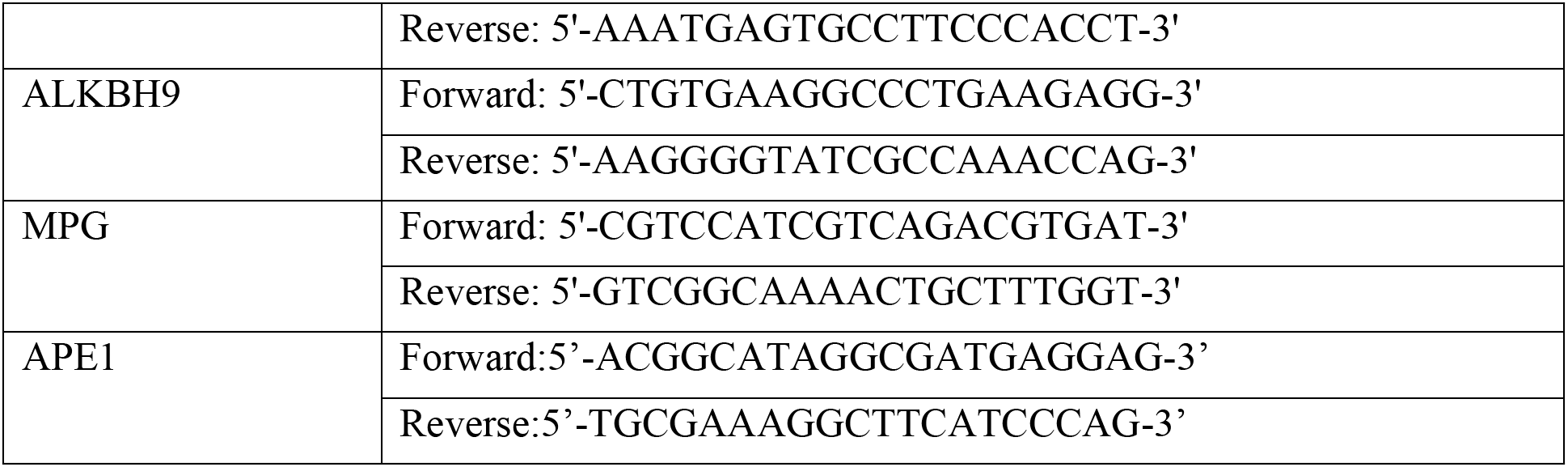
Sequences of qPCR primers.

**Supplementary Figure 1.**
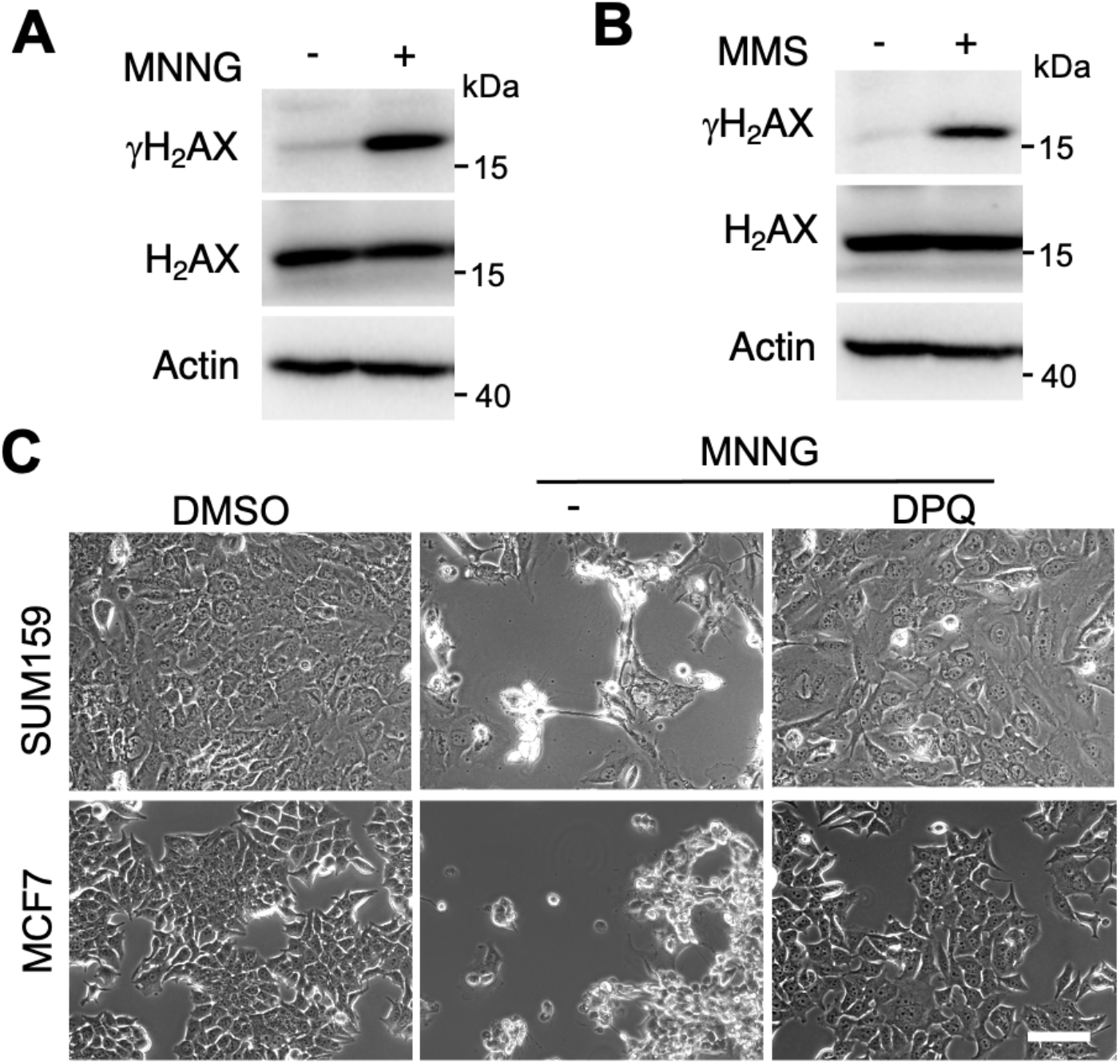
PARP-1 is required for alkylating agent MNNG-induced cell death in different types of cancer cells. (**A** and **B**) Immunoblot analysis of DNA damage in HeLa cells 6 h after the treatment with vehicle (−), MNNG (50 μM, 15 min), or MMS (2 mM, 6 h). (**C**) Representative cell death images from SUM159 and MCF-7 cells treated with DMSO, MNNG (200 μM for SUM159 and 500 μM for MCF7), MNNG plus DPQ (30 μM) for 24 h. Scale bar, 20 μm.

**Supplementary Figure 2.**
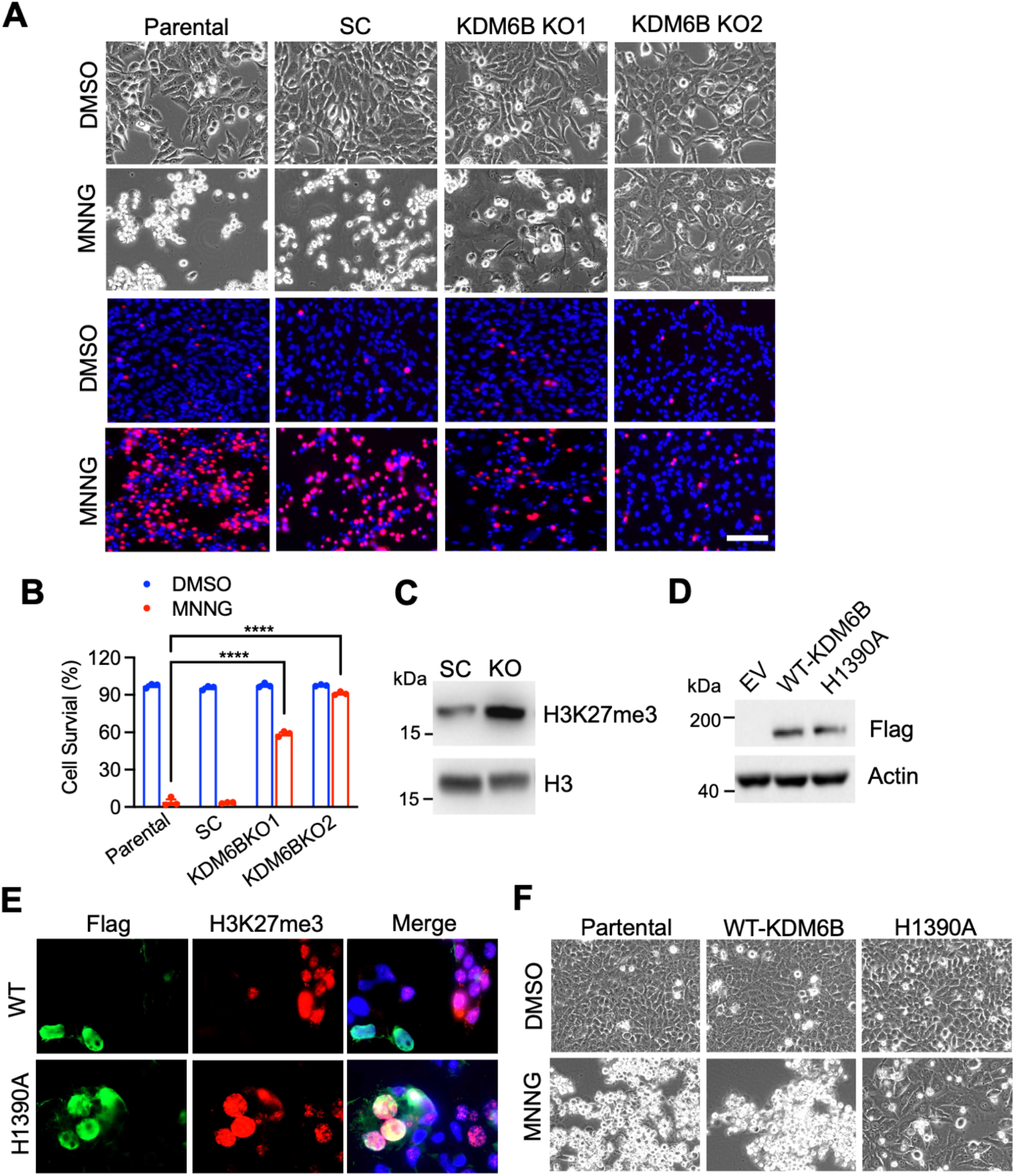
Loss of KDM6B or its demethylase activity blocked PARthanatos induced by alkylating agent MNNG. (**A** and **B**) Representative images and quantification of MNNG (50 μM, 15 min)-induced cell death in WT and KDM6B KO cells, which was established by sgRNAs targeting to different regions of *KDM6B* (mean ± SEM, *n* = 3). \*\*\*\**p* < 0.0001 by two-way ANOVA Sidak’s multiple comparisons test. (**C**) Expression of H3K27me3 in SC and KDM6B KO cells. (**D**) Expression of full-length Flag-tagged WT-KDM6B and demethylase inactive KDM6B H1390A mutant in HeLa cells. (**E**) Expression of H3K27me3 (red) and Flag-tagged WT and H1390A KDM6B (green) in HeLa cells overexpressed WT or H1390A mutant KDM6B. Blue indicates DAPI staining. (**F**) PARthanatos induced by MNNG in HeLa cells overexpressing WT or H1390A mutant KDM6B.

**Supplementary Figure 3.**
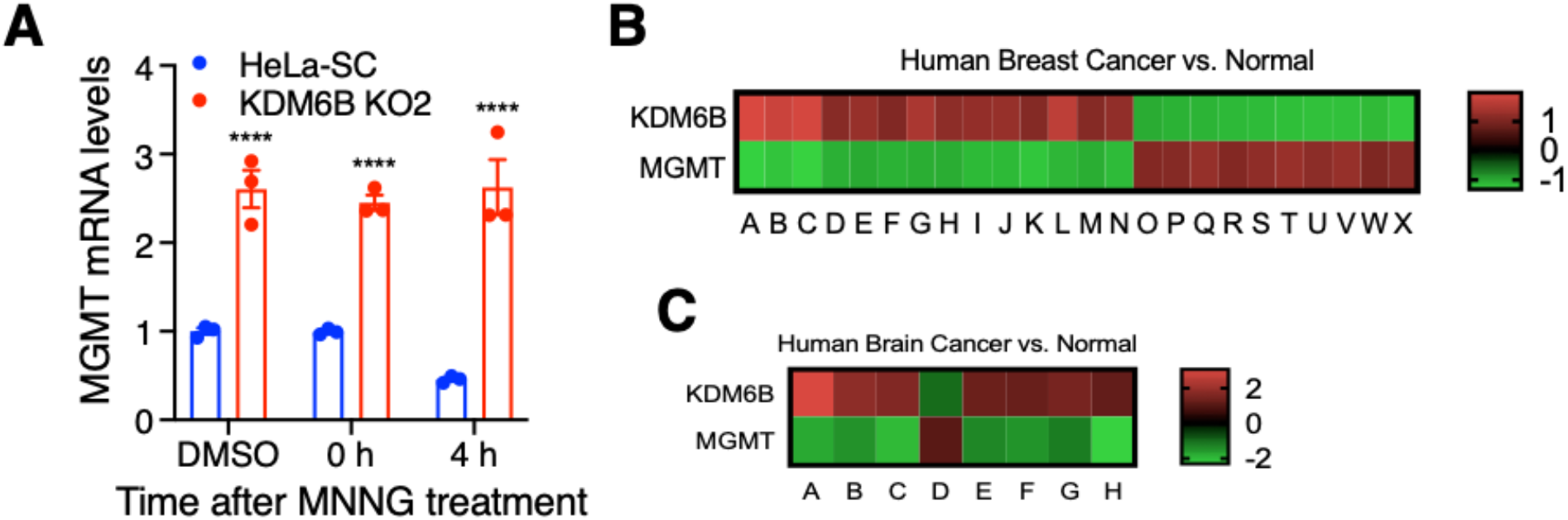
KDM6B negatively regulates MGMT expression. (**A**) RT-qPCR analysis of MGMT mRNA in SC and KDM6B KO2 HeLa cells treated with DMSO or MNNG for 15 min, followed by recovery for 0 and 4 h (mean ± SEM, *n* = 3). \*\*\*\**p* < 0.0001 by two-way ANOVA Sidak’s multiple comparisons test. (**B** and **C**) The heatmap reveals a negative correlation of KDM6B with MGMT mRNA in human breast (B) and brain (C) cancers. Data were retrieved from Oncomine datasets. The alphabet represents different data sets.

**Supplementary Figure 4.**
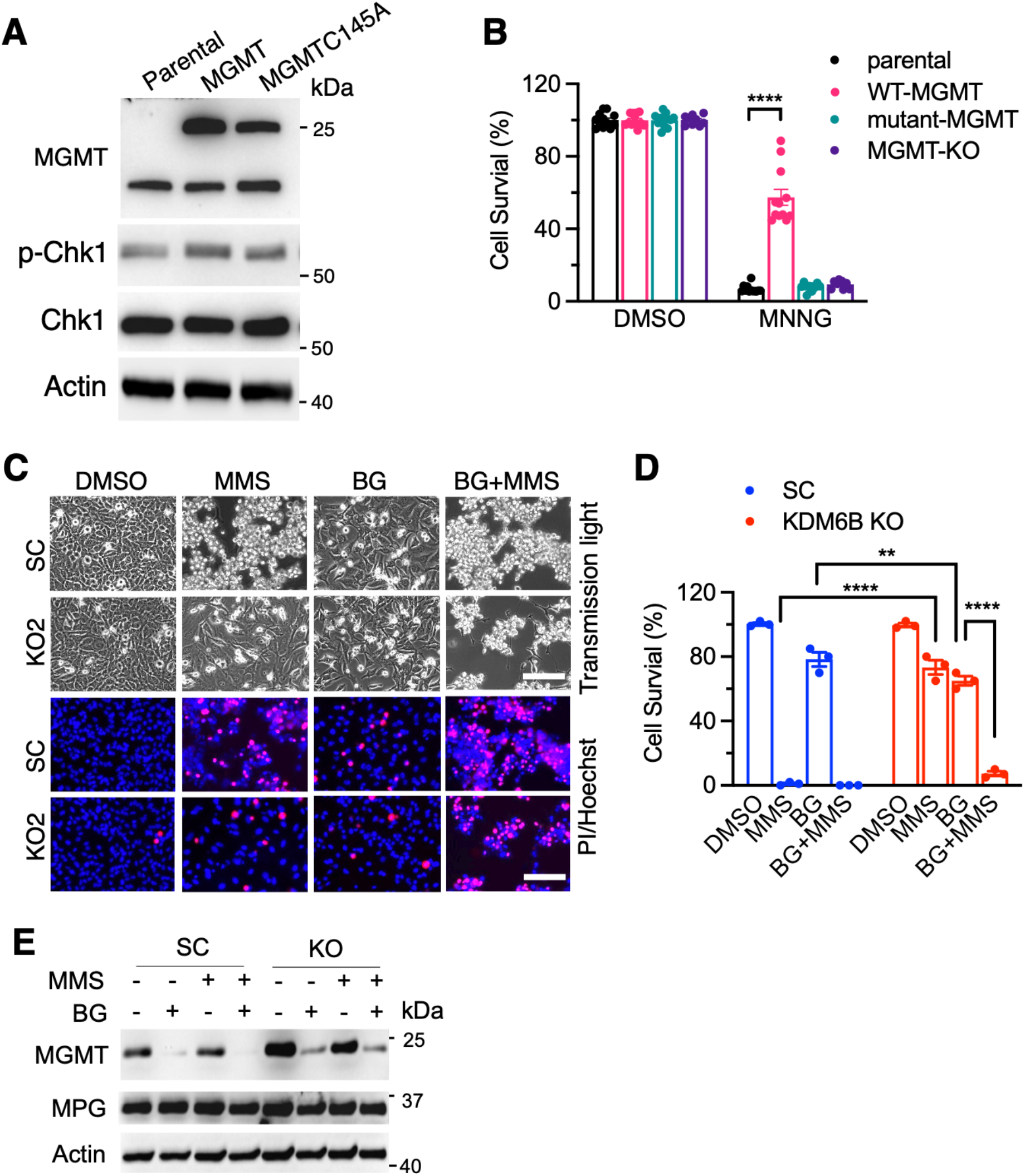
MGMT inhibition overcomes alkylating agent resistance in KDM6B KO cells. (**A**) Expression of MGMT and its methyltransferase inactive mutant C145A MGMT in HeLa cells. (**B**) Effects of MGMT, mutant C145A MGMT and MGMT KO on MNNG (50 μM, 15 min)-induced cell death (mean ± SEM, *n* = 9-12). \*\*\*\**p* < 0.0001 by two-way ANOVA Tukey’s multiple comparisons test. (**C** and **D**) Representative cell death images and quantification of cell death in SC and KDM6B KO2 HeLa cells 24 h after the treatment with MMS (2 mM, 1 h) and/or BG (200 μM) (mean ± SEM, *n* = 3). \*\**p* <0.01, \*\*\*\**p* < 0.0001 by two-way ANOVA Sidak’s multiple comparisons test. (**E**) Protein expression of MGMT and MPG 6 h after the treatment of MMS (2 mM, 1 h) and BG (200 μM).

## References

1. Ray Chaudhuri, A. and Nussenzweig, A. (2017) The multifaceted roles of PARP1 in DNA repair and chromatin remodelling. Nat Rev Mol Cell Biol, 18, 610–621.

2. Luo, X. and Kraus, W.L. (2012) On PAR with PARP: cellular stress signaling through poly(ADP-ribose) and PARP-1. Genes Dev, 26, 417–432.

3. Bai, P. (2015) Biology of Poly(ADP-Ribose) Polymerases: The Factotums of Cell Maintenance. Mol Cell, 58, 947–958.

4. Beck, C., Robert, I., Reina-San-Martin, B., Schreiber, V. and Dantzer, F. (2014) Poly(ADP-ribose) polymerases in double-strand break repair: focus on PARP1, PARP2 and PARP3. Exp Cell Res, 329, 18–25.

5. Wang, Y., Luo, W. and Wang, Y. (2019) PARP-1 and its associated nucleases in DNA damage response. DNA Repair (Amst), 81, 102651.

6. Bryant, H.E., Schultz, N., Thomas, H.D., Parker, K.M., Flower, D., Lopez, E., Kyle, S., Meuth, M., Curtin, N.J. and Helleday, T. (2005) Specific killing of BRCA2-deficient tumours with inhibitors of poly(ADP-ribose) polymerase. Nature, 434, 913–917.

7. Liu, S., Luo, W. and Wang, Y. (2021) Emerging role of PARP-1 and PARthanatos in ischemic stroke. J Neurochem.

8. Wang, Y., An, R., Umanah, G.K., Park, H., Nambiar, K., Eacker, S.M., Kim, B., Bao, L., Harraz, M.M., Chang, C. et al. (2016) A nuclease that mediates cell death induced by DNA damage and poly(ADP-ribose) polymerase-1. Science, 354.

9. Wang, Y., Kim, N.S., Haince, J.F., Kang, H.C., David, K.K., Andrabi, S.A., Poirier, G.G., Dawson, V.L. and Dawson, T.M. (2011) Poly(ADP-ribose) (PAR) binding to apoptosis-inducing factor is critical for PAR polymerase-1-dependent cell death (parthanatos). Sci Signal, 4, ra20.

10. Fu, D., Calvo, J.A. and Samson, L.D. (2012) Balancing repair and tolerance of DNA damage caused by alkylating agents. Nature reviews. Cancer, 12, 104–120.

11. Maanen, M.J., Smeets, C.J. and Beijnen, J.H. (2000) Chemistry, pharmacology and pharmacokinetics of N,N’,N” -triethylenethiophosphoramide (ThioTEPA). Cancer treatment reviews, 26, 257–268.

12. Lawley, P.D. and Thatcher, C.J. (1970) Methylation of deoxyribonucleic acid in cultured mammalian cells by N-methyl-N’-nitro-N-nitrosoguanidine. The influence of cellular thiol concentrations on the extent of methylation and the 6-oxygen atom of guanine as a site of methylation. Biochem J, 116, 693–707.

13. Zong, W.X., Ditsworth, D., Bauer, D.E., Wang, Z.Q. and Thompson, C.B. (2004) Alkylating DNA damage stimulates a regulated form of necrotic cell death. Genes Dev, 18, 1272–1282.

14. Fu, D., Jordan, J.J. and Samson, L.D. (2013) Human ALKBH7 is required for alkylation and oxidation-induced programmed necrosis. Genes Dev, 27, 1089–1100.

15. Klapacz, J., Meira, L.B., Luchetti, D.G., Calvo, J.A., Bronson, R.T., Edelmann, W. and Samson, L.D. (2009) O6-methylguanine-induced cell death involves exonuclease 1 as well as DNA mismatch recognition in vivo. Proc Natl Acad Sci U S A, 106, 576–581.

16. Calvo, J.A., Moroski-Erkul, C.A., Lake, A., Eichinger, L.W., Shah, D., Jhun, I., Limsirichai, P., Bronson, R.T., Christiani, D.C., Meira, L.B. et al. (2013) Aag DNA glycosylase promotes alkylation-induced tissue damage mediated by Parp1. PLoS Genet, 9, e1003413.

17. Chen, Y., Zhang, B., Bao, L., Jin, L., Yang, M., Peng, Y., Kumar, A., Wang, J.E., Wang, C., Zou, X. et al. (2018) ZMYND8 acetylation mediates HIF-dependent breast cancer progression and metastasis. J Clin Invest, 128, 1937–1955.

18. Zhang, B., Chen, Y., Shi, X., Zhou, M., Bao, L., Hatanpaa, K.J., Patel, T., DeBerardinis, R.J., Wang, Y. and Luo, W. (2021) Regulation of branched-chain amino acid metabolism by hypoxia-inducible factor in glioblastoma. Cell Mol Life Sci, 78, 195–206.

19. Shalem, O., Sanjana, N.E., Hartenian, E., Shi, X., Scott, D.A., Mikkelson, T., Heckl, D., Ebert, B.L., Root, D.E., Doench, J.G. et al. (2014) Genome-scale CRISPR-Cas9 knockout screening in human cells. Science, 343, 84–87.

20. Li, W., Xu, H., Xiao, T., Cong, L., Love, M.I., Zhang, F., Irizarry, R.A., Liu, J.S., Brown, M. and Liu, X.S. (2014) MAGeCK enables robust identification of essential genes from genome-scale CRISPR/Cas9 knockout screens. Genome Biol, 15, 554.

21. Hattermann, K., Mehdorn, H.M., Mentlein, R., Schultka, S. and Held-Feindt, J. (2008) A methylation-specific and SYBR-green-based quantitative polymerase chain reaction technique for O6-methylguanine DNA methyltransferase promoter methylation analysis. Anal Biochem, 377, 62–71.

22. De Santa, F., Totaro, M.G., Prosperini, E., Notarbartolo, S., Testa, G. and Natoli, G. (2007) The histone H3 lysine-27 demethylase Jmjd3 links inflammation to inhibition of polycomb-mediated gene silencing. Cell, 130, 1083–1094.

23. Yang, L., Zha, Y., Ding, J., Ye, B., Liu, M., Yan, C., Dong, Z., Cui, H. and Ding, H.F. (2019) Histone demethylase KDM6B has an anti-tumorigenic function in neuroblastoma by promoting differentiation. Oncogenesis, 8, 3.

24. Cabrini, G., Fabbri, E., Lo Nigro, C., Dechecchi, M.C. and Gambari, R. (2015) Regulation of expression of O6-methylguanine-DNA methyltransferase and the treatment of glioblastoma (Review). International journal of oncology, 47, 417–428.

25. Jin, J., Shirogane, T., Xu, L., Nalepa, G., Qin, J., Elledge, S.J. and Harper, J.W. (2003) SCFbeta-TRCP links Chk1 signaling to degradation of the Cdc25A protein phosphatase. Genes Dev, 17, 3062–3074.

26. Hazra, T.K., Roy, R., Biswas, T., Grabowski, D.T., Pegg, A.E. and Mitra, S. (1997) Specific recognition of O6-methylguanine in DNA by active site mutants of human O6-methylguanine-DNA methyltransferase. Biochemistry, 36, 5769–5776.

27. Li, M., Huang, T., Li, X., Shi, Z., Sheng, Y., Hu, M. and Song, K. (2021) GDC-0575, a CHK1 Inhibitor, Impairs the Development of Colitis and Colitis-Associated Cancer by Inhibiting CCR2(+) Macrophage Infiltration in Mice. Onco Targets Ther, 14, 2661–2672.

28. Chung, N., Bogliotti, Y.S., Ding, W., Vilarino, M., Takahashi, K., Chitwood, J.L., Schultz, R.M. and Ross, P.J. (2017) Active H3K27me3 demethylation by KDM6B is required for normal development of bovine preimplantation embryos. Epigenetics, 12, 1048–1056.

29. Chan, K.M., Fang, D., Gan, H., Hashizume, R., Yu, C., Schroeder, M., Gupta, N., Mueller, S., James, C.D., Jenkins, R. et al. (2013) The histone H3.3K27M mutation in pediatric glioma reprograms H3K27 methylation and gene expression. Genes Dev, 27, 985–990.

30. Rose, M., Burgess, J.T., O’Byrne, K., Richard, D.J. and Bolderson, E. (2020) PARP Inhibitors: Clinical Relevance, Mechanisms of Action and Tumor Resistance. Front Cell Dev Biol, 8, 564601.

31. Wang, Y., Dawson, V.L. and Dawson, T.M. (2009) Poly(ADP-ribose) signals to mitochondrial AIF: a key event in parthanatos. Exp Neurol, 218, 193–202.

